# Integrating hydrogen-exchange experiments with conformational ensembles

**DOI:** 10.64898/2026.11.23.768707

**Authors:** Julian O. Streit, Daniele Peterle, Giulio Tesei, Michele Invernizzi, Sandro Bottaro, Thea K. Schulze, Patrick Kunz, Kamil Tamiola, Kresten Lindorff-Larsen

## Abstract

Protein conformational dynamics are central to biological function but remain difficult to characterise at scale. Amide hydrogen exchange (HX), measured by NMR or mass spectrometry, probes these dynamics but requires computational models for integration with structural ensembles. We benchmark models for calculating HX from ensembles using a large literature-derived dataset spanning folded and intrinsically disordered proteins. We then develop a model that identifies exchange-competent amides from local structural descriptors while accounting for electrostatic effects. The model predicts protection across a broad dynamic range in ensembles generated by molecular dynamics and generative AI. HX-guided ensemble reweighting improves agreement with independent NMR, SAXS, and folding-stability measurements. Millisecond HX distinguishes and refines alternative ensembles of intrinsically disordered proteins, while reweighting of structured proteins reveals folding intermediates and low-population excited states. These results establish HX as a quantitative restraint for ensemble reweighting and provide a scalable framework for benchmarking simulations and generative protein models.

## Introduction

The success of AlphaFold2 and related structure-prediction methods has transformed protein structural biology by making accurate structural models available at unprecedented scale [1–3]. A major next frontier is to move beyond single structures towards the prediction and validation of conformational dynamics, structural ensembles and low-population states in their biological context [4, 5]. The functional behaviour of intrinsically disordered proteins (IDPs) [6], proteins during folding and aggregation processes [7, 8], and allosteric protein-ligand interactions [9, 10] can all depend on transiently populated conformations. Progress therefore requires not only improved molecular simulations and generative models, but also experimental measurements that can quantitatively evaluate and refine conformational ensembles at scale [5, 11].

Amide hydrogen exchange (HX) provides a sensitive probe of conformational ensembles and protein dynamics [12–14]. Amide hydrogens exchange with solvent at rates determined by their intrinsic chemical reactivity and by structural fluctuations that transiently render them exchange competent [12, 15, 16]. Under EX2 conditions, the observed exchange rate reflects the equilibrium population of exchange-competent conformations, such that the corresponding protection factor reports on the free-energy difference associated with structural opening [12, 13]. HX can therefore report on persistent structure, transient local unfolding, fluctuations in solvent accessibility and low-population conformational states over a broad range of timescales [13].

HX can be measured at residue resolution by NMR or through hydrogen-deuterium exchange mass spectrometry (HDX-MS), which typically reports deuterium uptake at peptide resolution (Figure 1a). HDX-MS is particularly attractive because it can be applied to large, dynamic proteins and macromolecular complexes [10, 17, 18], complementing high-resolution methods for protein structure determination [18, 19]. It is also well suited to measurements across many sequences and experimental conditions at comparatively high throughput [14, 20]. These properties make HDX-MS a promising quantitative complement to other structural techniques and, potentially, a scalable source of experimental data for benchmarking and ultimately training molecular simulation force fields and generative protein models. Realising this potential, however, requires accurate and transferable ‘forward models’ that are used to calculate the measured exchange observables from structural ensembles (Figure 1a).

**Fig. 1.**
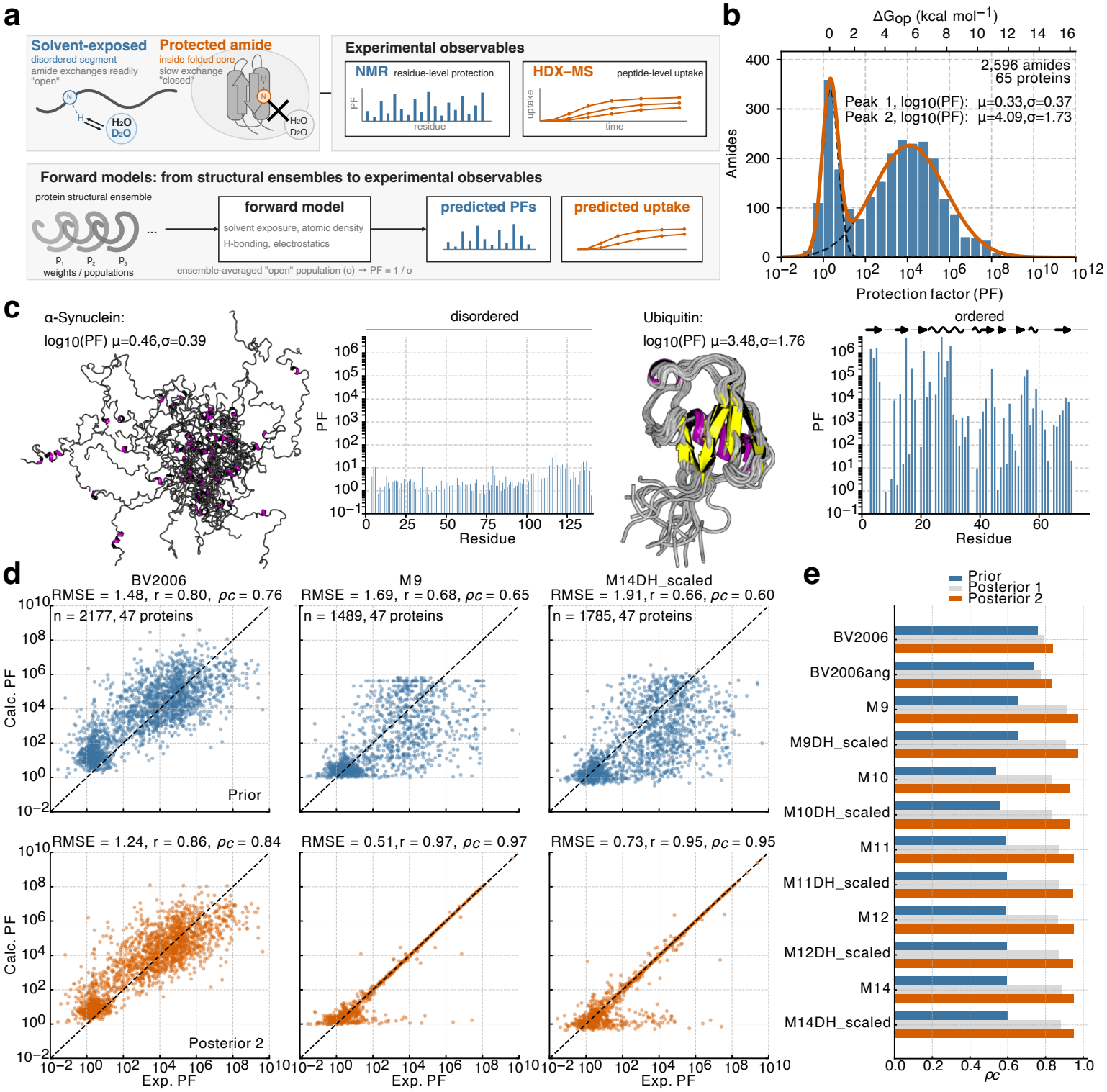
HX-NMR dataset and comparison of forward models before and after ensemble reweighting using HX protection factors. **a**. Schematic of amide hydrogen exchange (HX) with solvent protons or deuterons. Solvent-exposed amides exchange readily, whereas buried or hydrogen-bonded amides are protected from exchange. HX can be measured experimentally by NMR, yielding residue-resolved protection factors (PFs), or by HDX–MS, yielding peptide-resolved deuterium uptake curves. Forward models predict these experimental observables from structural ensembles based on properties such as solvent exposure, atomic density, hydrogen bonding, and electrostatics. **b**. Distribution of experimental PFs in the HX-NMR dataset, comprising 2,596 amides in 65 proteins. PFs are shown on a logarithmic scale, with the corresponding opening free energy (Δ*G*_op_) indicated on the upper x-axis. Two Gaussian components were fitted to the log_10_(PF) distribution to characterise their means (*µ*) and standard deviations (*σ*). **c**. Representative PF profiles for the intrinsically disordered protein *α*-synuclein and the folded protein ubiquitin. *α*-synuclein exhibits uniformly low PFs, close to unity, consistent with its lack of persistent secondary and tertiary structure. In contrast, ubiquitin exhibits heterogeneous and substantially higher PFs spanning several orders of magnitude, characteristic of a folded protein with regions of strong protection. **d**. Correlation between experimental (Exp.) and calculated (Calc.) PFs across amides and proteins in the dataset for three representative forward models (BV2006, M9, and M14DH_scaled). Predictions are shown for the prior ensembles (top) and after ensemble reweighting (posterior 2; bottom). Agreement with experimental PFs was quantified in log_10_ space by the root-mean-square error (RMSE), Pearson correlation coefficient (*r*), and concordance correlation coefficient (*ρ*_*c*_). All 2,177 amides were included for the empirical model, BV2006, whereas analyses using explicit models were restricted to amides that sampled an open state at least once under the respective model definition. **e**. Performance of the tested forward models, quantified by *ρ*_*c*_, for the prior ensembles and after reweighting. posterior 1 corresponds to the elbow of the optimization L-curve, whereas posterior 2 corresponds to a minimum fraction of contributing frames of *ϕ* = 0.1.

Existing forward models that connect structures to HX can broadly be divided into two classes. Empirical models typically estimate protection directly from structural descriptors of the ground-state ensemble, using combinations of hydrogen bonding, solvent accessibility, local packing or contact measures [21–23]. In other words, they have been devised to predict opening free-energies from structures where amides are predominantly protected from exchange (i.e., folded regions). While these models have been applied extensively to shed light on protein dynamics [24–27], their predictive accuracy and transferability across diverse proteins and conformational regimes remains unclear, particularly for disordered and partially unfolded states. A second class of approaches attempts to model exchange more mechanistically by explicitly identifying exchange-competent, or open, conformations from simulation [23, 28, 29]. Such models avoid assigning protection solely from the closed state, but have so far been applied to relatively few systems and remain only sparsely validated against independent experimental data. More generally, rigorous validation of either class is difficult because experimentally determined ground-truth conformational ensembles are rarely available. Agreement with HX observables alone therefore does not establish that the underlying structural populations are correct.

Here we address these challenges by assembling a large literature-derived HX dataset spanning dynamics within folded and intrinsically disordered proteins and integrating it with molecular simulations, generative ensembles, and complementary NMR, SAXS and folding-stability measurements. This framework enables systematic evaluation of HX forward models, Bayesian/maximum-entropy ensemble reweighting [30–32], and cross-validation against independent experimental observables. We develop a transferable residue-level HX model in which exchange competence is defined explicitly from local structural descriptors and modulated by electrostatic effects, thereby linking measured protection directly to the structures and populations of exchange-competent conformations. Combined with state-of-the-art millisecond HDX-MS measurements of intrinsically disordered proteins, this approach enables alternative conformational ensembles to be distinguished by their agreement with experiment and quantitatively refined by reweighting. Applied to folded proteins, it further reveals and quantifies folding intermediates and low-population excited states. Together, these advances establish HX as a quantitative method for ensemble refinement and position HDX-MS as a scalable source of structural information for benchmarking, refining and ultimately training the next generation of molecular simulation force fields and generative protein models. Our forward models and reweighting methods are implemented in an open-source Python package, HXEnsemble (https://github.com/KULL-Centre/HXEnsemble).

## Results

### A dataset of HX-NMR protection factors and conformational ensembles

To enable systematic evaluation of forward models, we assembled an extensive literature-derived dataset of HX-NMR (where HX denotes amide hydrogen exchange in general, whereas HDX refers specifically to amide hydrogen–deuterium exchange) protection factors (PFs) covering 65 proteins (Methods; Table 1). PFs report on the extent to which an amide is protected against exchange, and in the EX2 regime of the Linderstrøm-Lang model for exchange, the logarithm of the PF reports on the free-energy difference between the ground (closed) and exchange-competent (open) state. The dataset spans both folded and intrinsically disordered proteins and exhibits a bimodal distribution of PFs (Figure 1b), with one population centred at log_10_ PF ~ 0.3, characteristic of weakly protected disordered regions, and a second centred at log_10_ PF = 4.1, characteristic of strongly protected folded regions. Overall, the measurements span approximately ten orders of magnitude in PF, corresponding to local opening free energies (Δ*G*_op_) of up to ~12 kcal mol^*−*1^. As representative examples, *α*-synuclein displays uniformly low protection across most of its sequence, with PFs predominantly between 1 and 10 [33], whereas ubiquitin exhibits substantially stronger and more heterogeneous protection spanning several orders of magnitude [34] (Figure 1c).

**Table 1:**
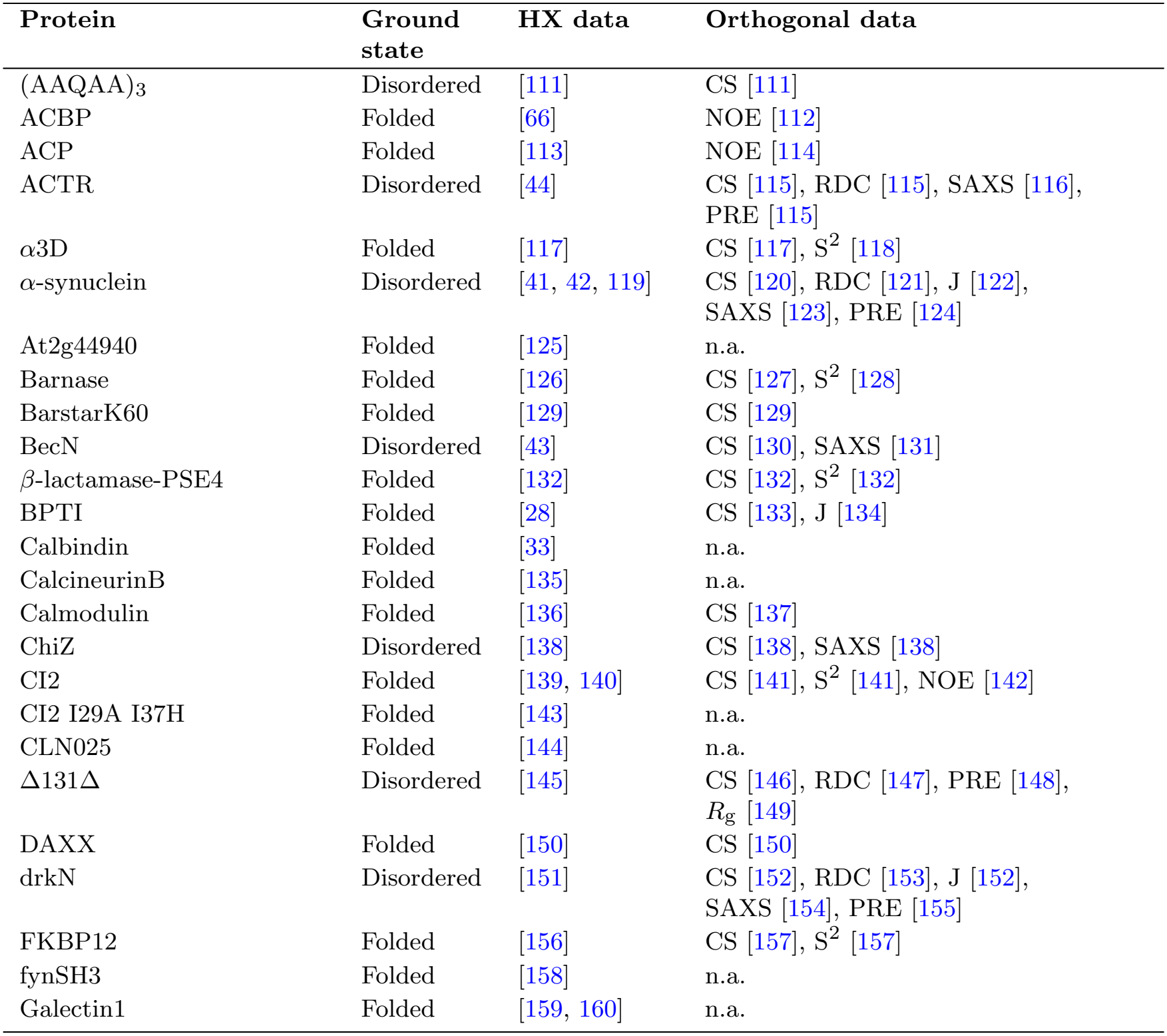
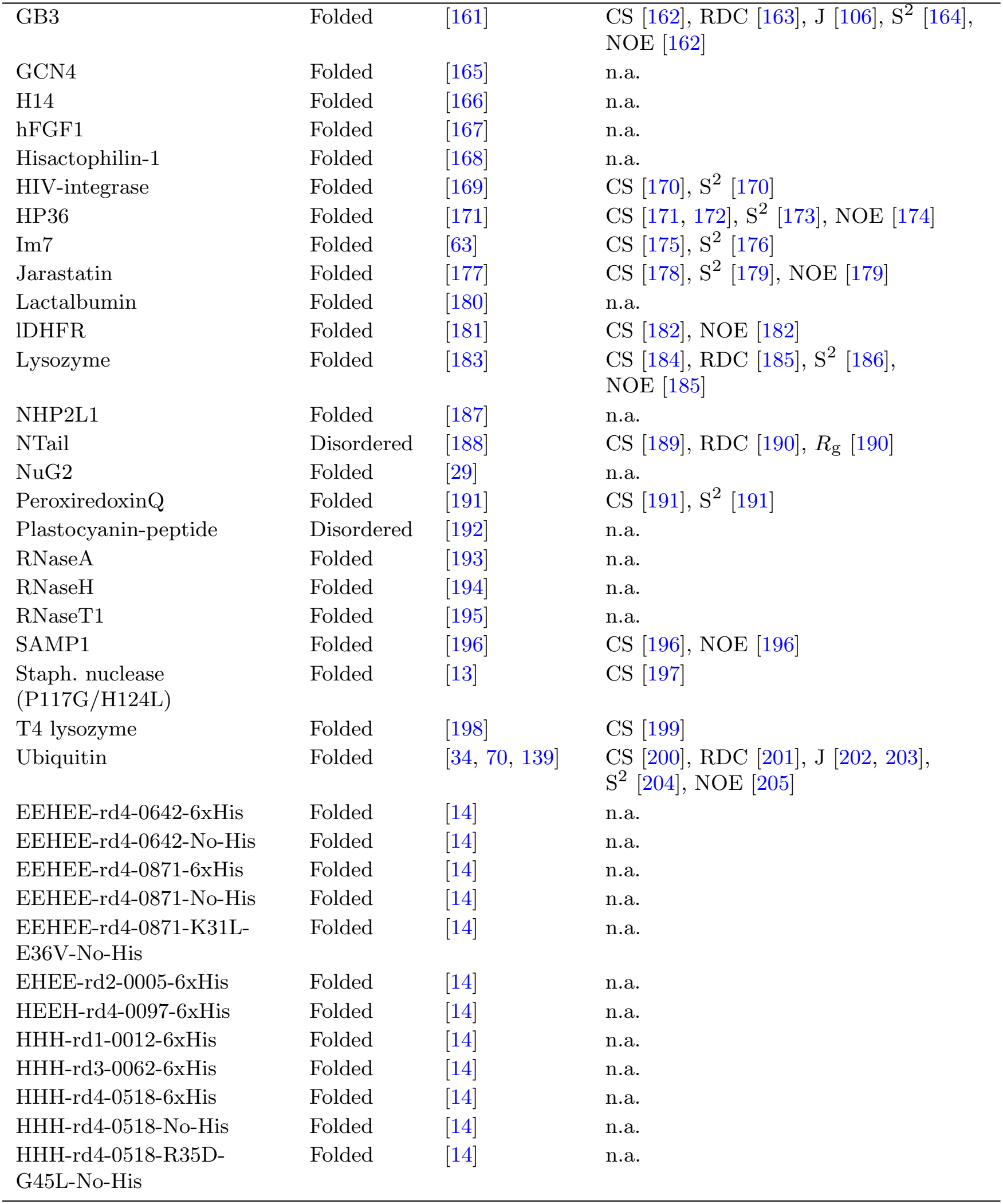
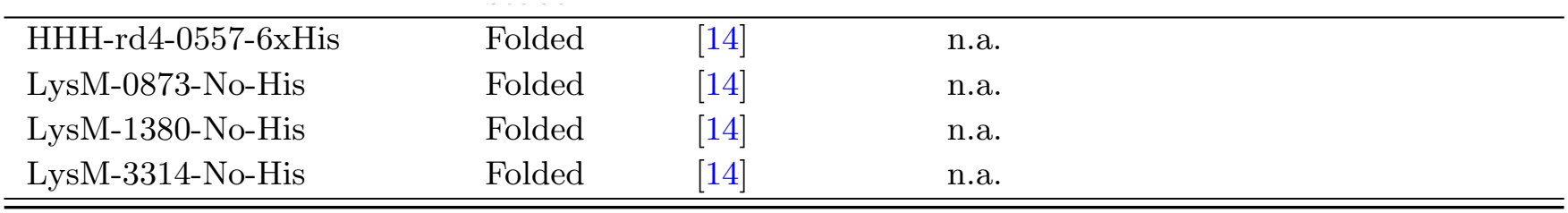
HX-NMR literature dataset and corresponding orthogonal experimental data reporting on structural and dynamic properties of the ground state. Some datasets measure H–H or D–H exchange rather than H–D exchange; for simplicity, all measurements are referred to collectively as hydrogen exchange (HX). CS, backbone chemical shifts; RDC, amide residual dipolar couplings; J, backbone ^3^J-couplings; PRE, paramagnetic relaxation enhancement; SAXS, small-angle X-ray scattering; *R*_g_, radius of gyration; NOE, nuclear Overhauser effect distances; S^2^, amide order parameters; n.a., not applicable.

| Protein | Ground state | HX data | Orthogonal data |
|---|---|---|---|
| (AAQAA)<sub>3</sub> | Disordered | [111] | CS [111] |
| ACBP | Folded | [66] | NOE [112] |
| ACP | Folded | [113] | NOE [114] |
| ACTR | Disordered | [44] | CS [115], RDC [115], SAXS [116], PRE [115] |
| α3D | Folded | [117] | CS [117], S<sup>2</sup> [118] |
| α-synuclein | Disordered | [41, 42, 119] | CS [120], RDC [121], J [122], SAXS [123], PRE [124] |
| At2g44940 | Folded | [125] | n.a. |
| Barnase | Folded | [126] | CS [127], S<sup>2</sup> [128] |
| BarstarK60 | Folded | [129] | CS [129] |
| BecN | Disordered | [43] | CS [130], SAXS [131] |
| β-lactamase-PSE4 | Folded | [132] | CS [132], S<sup>2</sup> [132] |
| BPTI | Folded | [28] | CS [133], J [134] |
| Calbindin | Folded | [33] | n.a. |
| CalcineurinB | Folded | [135] | n.a. |
| Calmodulin | Folded | [136] | CS [137] |
| ChiZ | Disordered | [138] | CS [138], SAXS [138] |
| CI2 | Folded | [139, 140] | CS [141], S<sup>2</sup> [141], NOE [142] |
| CI2 I29A I37H | Folded | [143] | n.a. |
| CLN025 | Folded | [144] | n.a. |
| Δ131Δ | Disordered | [145] | CS [146], RDC [147], PRE [148], *R*<sub>g</sub> [149] |
| DAXX | Folded | [150] | CS [150] |
| drkN | Disordered | [151] | CS [152], RDC [153], J [152], SAXS [154], PRE [155] |
| FKBP12 | Folded | [156] | CS [157], S<sup>2</sup> [157] |
| fynSH3 | Folded | [158] | n.a. |
| Galectin1 | Folded | [159, 160] | n.a. |
| GB3 | Folded | [161] | CS [162], RDC [163], J [106], S<sup>2</sup> [164], NOE [162] |
| GCN4 | Folded | [165] | n.a. |
| H14 | Folded | [166] | n.a. |
| hFGF1 | Folded | [167] | n.a. |
| Hisactophilin-1 | Folded | [168] | n.a. |
| HIV-integrase | Folded | [169] | CS [170], S<sup>2</sup> [170] |
| HP36 | Folded | [171] | CS [171, 172], S<sup>2</sup> [173], NOE [174] |
| Im7 | Folded | [63] | CS [175], S<sup>2</sup> [176] |
| Jarastatin | Folded | [177] | CS [178], S<sup>2</sup> [179], NOE [179] |
| Lactalbumin | Folded | [180] | n.a. |
| lDHFR | Folded | [181] | CS [182], NOE [182] |
| Lysozyme | Folded | [183] | CS [184], RDC [185], S<sup>2</sup> [186], NOE [185] |
| NHP2L1 | Folded | [187] | n.a. |
| NTail | Disordered | [188] | CS [189], RDC [190], *R*<sub>g</sub> [190] |
| NuG2 | Folded | [29] | n.a. |
| PeroxiredoxinQ | Folded | [191] | CS [191], S<sup>2</sup> [191] |
| Plastocyanin-peptide | Disordered | [192] | n.a. |
| RNaseA | Folded | [193] | n.a. |
| RNaseH | Folded | [194] | n.a. |
| RNaseT1 | Folded | [195] | n.a. |
| SAMP1 | Folded | [196] | CS [196], NOE [196] |
| Staph. nuclease (P117G/H124L) | Folded | [13] | CS [197] |
| T4 lysozyme | Folded | [198] | CS [199] |
| Ubiquitin | Folded | [34, 70, 139] | CS [200], RDC [201], J [202, 203], S<sup>2</sup> [204], NOE [205] |
| EEHEE-rd4-0642-6xHis | Folded | [14] | n.a. |
| EEHEE-rd4-0642-No-His | Folded | [14] | n.a. |
| EEHEE-rd4-0871-6xHis | Folded | [14] | n.a. |
| EEHEE-rd4-0871-No-His | Folded | [14] | n.a. |
| EEHEE-rd4-0871-K31L-E36V-No-His | Folded | [14] | n.a. |
| EHEE-rd2-0005-6xHis | Folded | [14] | n.a. |
| HEEH-rd4-0097-6xHis | Folded | [14] | n.a. |
| HHH-rd1-0012-6xHis | Folded | [14] | n.a. |
| HHH-rd3-0062-6xHis | Folded | [14] | n.a. |
| HHH-rd4-0518-6xHis | Folded | [14] | n.a. |
| HHH-rd4-0518-No-His | Folded | [14] | n.a. |
| HHH-rd4-0518-R35D-G45L-No-His | Folded | [14] | n.a. |
| HHH-rd4-0557-6xHis | Folded | [14] | n.a. |
| LysM-0873-No-His | Folded | [14] | n.a. |
| LysM-1380-No-His | Folded | [14] | n.a. |
| LysM-3314-No-His | Folded | [14] | n.a. |

For a subset of 47 proteins, we next assembled conformational ensembles by performing all-atom molecular dynamics (MD) simulations, totalling approximately 1.7 ms of sampling across the dataset (Methods; Table 3), together with previously published all-atom MD datasets (Table 4) and hybrid-resolution simulations performed with the HyRes-Opt force field [35] (Table 5). Together, these simulations provided ground-state conformational ensembles for both folded and disordered proteins in the HX-NMR dataset, allowing alternative HX forward models to be evaluated across a broad range of structural environments under the assumption that conformational dynamics are fast relative to intrinsic exchange, *k*_op_ + *k*_cl_ ≫ *k*_int_ (Methods).

### Forward models

A number of empirical forward models have been proposed to relate structural descriptors to amide protection factors. These commonly express ln *PF* as a function of ensemble-averaged quantities such as hydrogen bonding, solvent exposure, local packing, heavy-atom contacts, distances between the amide proton and polar atoms, or backbone RMSF [21–23, 36, 37]. Related approaches have used hydrogen-bond occupancies to protein and water molecules as predictors of protection [38, 39]. The premise of such models is that hydrogens that are buried or hydrogen-bonded in the ground state will have a large free-energy of opening, so that these features enable prediction of the PF from these ground-state ensembles. In contrast, explicit open-state models classify each amide in each conformation as either exchange competent or exchange incompetent, and predict the protection factor from the ensemble population of the exchange-competent state [23, 28]. Proposed criteria for defining this state include coordination by explicitly represented water molecules, solvent-accessible surface area (SASA) of the amide proton and the absence of amide hydrogen bonds [23, 28, 29].

In explicit open-state models, the protection factor of amide *i* is calculated from its ensemble-averaged effective exchange-competent population,

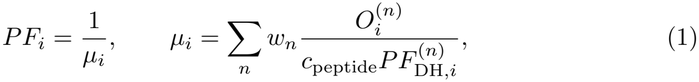

where *w*_*n*_ is the statistical weight of conformation *n*, 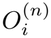 is a binary indicator equal to 1 when the amide is structurally open and 0 when closed, *c*_peptide_ is a peptide-based correction factor, and 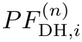 accounts for electrostatic modulation of exchange (Methods). Without these corrections, the protection factor reduces to the inverse of the ensemble-averaged open-state population. The explicit open-state models we analysed share this functional form but differ in the structural criteria used to classify amides as exchange competent.

We focused on two empirical solute-only models: the widely used Best-Vendruscolo model (BV2006), which—building on earlier work [21]—combines hydrogen-bonding and heavy-atom contact descriptors [22], and a modified variant, BV2006ang, that uses a more restrictive heavy-atom contact-distance cutoff and additionally imposes an angular criterion on hydrogen-bond assignment (Methods). We compared these with explicit open-state models M9–M12 and M14 (Methods). M9 classifies amides using amide-proton SASA, with an opening threshold of 4.0 Å^2^, together with the distance to nearby polar atoms, with an opening threshold of 3.0 Å and no angular criterion [23]. M10–M12 instead use an explicit geometric definition of hydrogen bonding together with a substantially lower SASA cutoff of 0.05 Å^2^, designed to distinguish complete from partial burial. M11 and M12 additionally incorporate alternative variants of the packing descriptor *ρ*_front_, which quantifies directional atomic density in front of the amide N–H bond. M14 uses a somewhat less stringent SASA cutoff of 0.5 Å^2^ together with *ρ*_front_, retaining directional steric shielding while allowing greater solvent exposure than M10–M12.

Under most experimental conditions, HX is predominantly a base-catalyzed reaction that depends on a negatively-charged hydroxide ion extracting the amide proton [16]. We therefore further introduced electrostatically corrected variants in which the local electrostatic environment modulates the effective exchange rate of structurally open amides [40, 41]. We denote these models MXDH and MXDH_scaled, where MX refers to the corresponding open-state model and DH_scaled uses uniformly scaled partial charges to approximately account for electronic polarisation effects using a Debye-Hückel model [41]. Unless otherwise stated, the main text analyses show the DH_scaled variants, while comparisons of the uncorrected, DH and DH_scaled models are provided in the Supplementary Information. Together, these models allowed us to test the relative contributions of hydrogen bonding, solvent exposure, directional steric shielding and electrostatics to a transferable description of HX protection.

Based on the benchmarks presented below, we use M14DH_scaled as our default model for subsequent applications, while retaining M11 and M12 for comparisons of alternative structural criteria.

### Explicit open-state models result in improved agreement after ensemble reweighting

We next compared predicted and experimental HX-NMR protection factors across 47 proteins for which corresponding ground-state MD ensembles were available. All ensembles were generated using all-atom MD simulations, except those of the IDPs *α*-synuclein, BecN, ChiZ and Δ131Δ, which were simulated using the HyRes-Opt force field (Tables 3– 5). We assessed each forward model both before and after Bayesian/maximum-entropy reweighting (Methods). Performance on the prior ensemble reflects how well a given model predicts protection from the unrefined ground-state ensemble. Post-reweighting agreement instead reflects how well the combination of forward model and sampled conformational ensemble can reproduce the experimental data when the prior deviates from the unknown ground truth. We considered two posterior ensembles: posterior 1, selected at the elbow of the reweighting L-curve and therefore remaining relatively close to the prior, and posterior 2, representing a more extensively reweighted solution constrained to retain an effective fraction of frames of *ϕ >* 0.1 (Supplementary Figure 1a). Of the 2,177 amides in this dataset, not all sampled an exchange-competent conformation under the different explicit model definitions (ranging from 1,489 for M9 to 1,785 for M14). For these models, reweighting and performance evaluation were therefore restricted to amides that sampled an open state at least once, whereas all amides were included for the empirical BV-based models.

Before reweighting, BV2006 showed the highest agreement with experiment among the tested models (*ρ*_*c*_ = 0.76, RMSE = 1.48 in log_10_ *PF* space; Figure 1d–e, Supplementary Figure 1b–c). This is consistent with its original development as an empirical predictor of protection from native-state structures [21, 22]. Re-optimising the BV2006 coefficients against our dataset yielded parameters close to the original values and only marginally improved agreement (Supplementary Figure 2a). BV2006 also out-performed the additional empirical M8 model based on SASA and distances from the amide proton to nearby polar atoms (Supplementary Figure 2c).

After reweighting, however, all tested explicit open-state models (M9–M12 and M14) outperformed the empirical BV2006 and BV2006ang models for both posterior 1 and posterior 2 (Figure 1d–e, Supplementary Figure 1b–c). Thus, while the empirical models provide accurate predictions of residues in folded regions based on ground-state ensembles, they are less amenable to further refinement by ensemble reweighting. The previously proposed M9 model and our M14DH_scaled model reached concordance correlation coefficients of *ρ*_*c*_ = 0.95–0.97, indicating that explicit treatment of exchange competence provides a substantially more effective basis for HX-guided ensemble refinement than empirical protection models.

We also evaluated the Persson–Halle (PH) explicit-water model [28] on peptide systems for which we generated explicit-solvent trajectories. Although much smaller than folded proteins, these systems provide a sensitive test of forward models because they represent weakly protected, highly dynamic states in which exchange should approach the intrinsic rate. In particular, short unstructured peptides and capped tripeptides offer a stringent baseline: in the absence of persistent secondary or tertiary structure, the model should predict little or no protection, with *PF* values close to unity (because exchange from such peptides is used to define the experimental baseline of no protection). Correctly reproducing this limit is important because systematic overprediction of protection in such systems would compress the dynamic range of the model and compromise its transferability to disordered and partially unfolded states. Across these systems, the PH model overpredicted protection by approximately one to two orders of magnitude (Supplementary Figure 3). M9 similarly showed a modest tendency to overpredict protection for model tripeptides, whereas M10–M12 and M14 reproduced the expected baseline of *PF* = 1 for short peptides lacking stable secondary or tertiary structure (Supplementary Figure 3). This demonstrates that the explicit open-state functional form alone does not guarantee accurate HX predictions; the structural criteria used to identify exchange-competent conformations are also critical. We therefore focused subsequent analyses on these explicit open-state models, with M14DH_scaled used as the principal model in the main text.

### Predicting protection factors from refined disordered-protein ensembles

We next evaluated forward-model performance using conformational ensembles expected to be closer to the experimental ground truth. For this purpose, we considered eight IDPs for which independent NMR and/or SAXS data were available. Fast amide exchange in IDPs is typically accessible by NMR measurements of H–H solvent exchange, rather than H–D exchange as commonly used for more strongly protected folded proteins, using experiments such as phase-modulated clean chemical exchange (CLEANEX-PM) [42, 43] or decorrelation by exchange (DECOR) [33, 44]. IDPs provide a useful setting for testing HX forward models because their protection generally arises from transient structural features rather than from extremely rare opening events. NMR chemical shifts and SAXS measurements are sensitive to ensemble-averaged secondary-structure propensity and global compaction, respectively, and can therefore be used to refine the same underlying conformational ensemble that gives rise to fast HX. We reasoned that IDP ensembles independently refined against such data should thus provide a more stringent test of forward-model accuracy than unrefined simulations alone. This comparison is less straightforward for folded proteins, where PFs can be several orders of magnitude larger and may be dominated by very low-population excited or partially unfolded states. Such rare conformations can contribute strongly to HX while having little influence on ensemble-averaged observables such as chemical shifts.

We refined ensembles for five IDPs simultaneously against NMR chemical shifts and SAXS data (ACTR, *α*-synuclein, BecN, drkN and ChiZ), and for three additional systems against NMR chemical shifts alone ((AAQAA)_3_, Δ131Δ and NTail, Methods; Tables 3– 5). Where available, the resulting ensembles were cross-validated against orthogonal observables not used during refinement, including residual dipolar couplings (RDCs) and paramagnetic relaxation enhancement (PRE) intensity ratios. As expected, reweighting improved agreement with the experimental observables used as restraints, but importantly also improved agreement with independent validation data where available, supporting an overall improvement in ensemble quality after refinement (Supplementary Figure 4a).

Using these refined ensembles, we next compared the ability of the different HX forward models to predict experimental protection factors. The empirical BV2006 and BV2006ang models systematically overpredicted protection, in some cases by up to an order of magnitude (Figure 2a, Supplementary Figure 4b–c), consistent with their original development using predominantly folded proteins. In contrast, the explicit open-state models showed substantially improved agreement when electrostatic effects were included, in line with previous work [41]. Across the tested models, addition of the electrostatic correction (Methods) increased *ρ*_*c*_, whereas an electrostatics-only model lacking structural descriptors such as SASA, hydrogen bonding and directional packing performed worse than models combining structural and electrostatic terms. The charge-scaled variants of M11, M12 and M14 performed best overall, with M14DH_scaled reaching *ρ*_*c*_ = 0.42 (Figure 2a–d, Supplementary Figure 4b–d). The same relative trend between forward models was already apparent before NMR/SAXS refinement, while independent refinement against these observables modestly improved the subsequent agreement with HX-NMR PFs (Supplementary Figure 4c). This provides further support that the refined ensembles more accurately capture the conformational features relevant to HX. Together, these results indicate that accurate prediction of HX protection in disordered proteins requires both stringent structural criteria for exchange competence and an explicit treatment of local electrostatic effects. The different agreement with experimental HX data among models sharing the same explicit open-state formulation also again highlights the importance of adequately defining exchange competence, particularly for the transiently populated structures characteristic of disordered proteins.

**Fig. 2.**
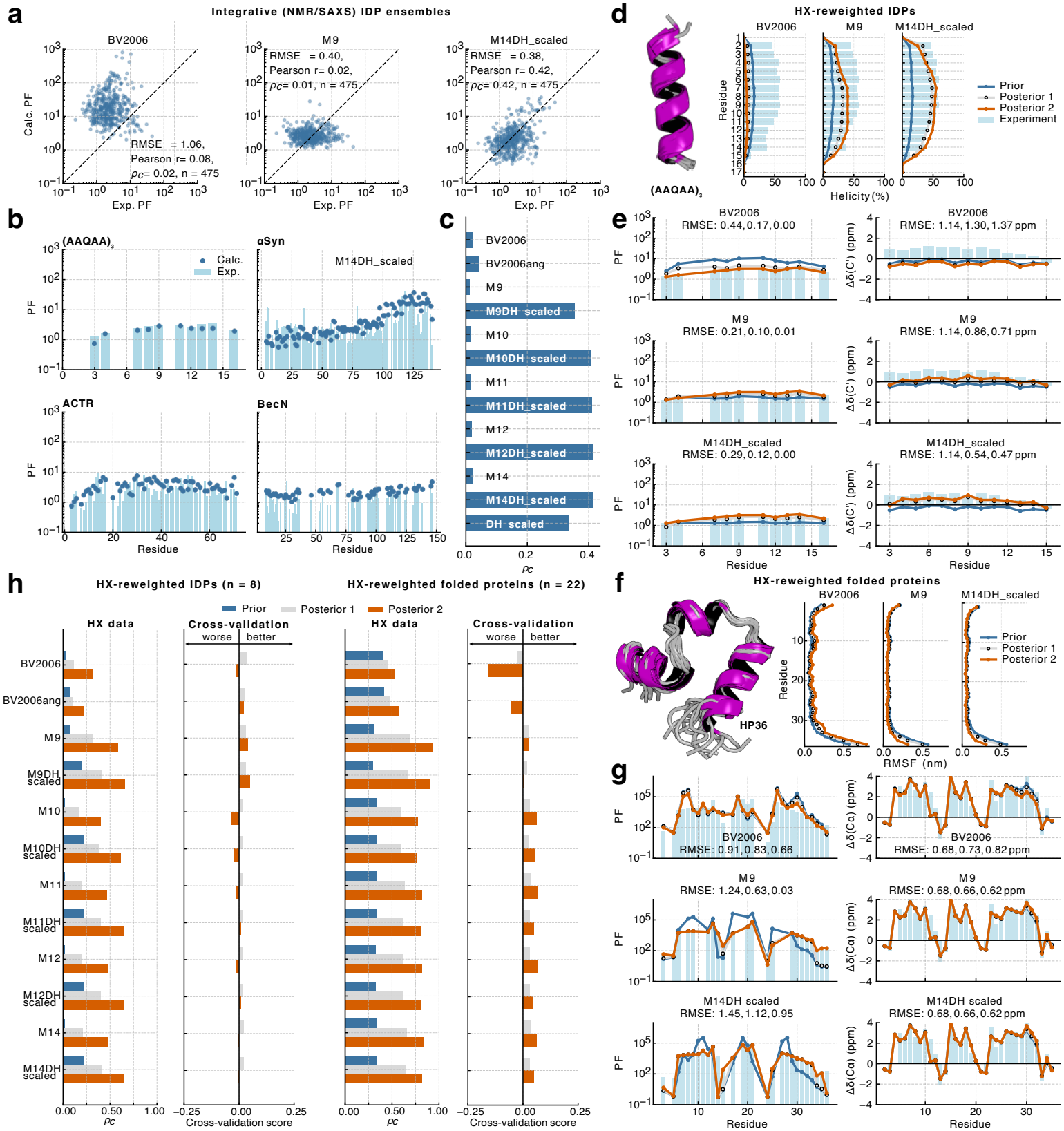
Benchmarking forward models on molecular dynamics ensembles of disordered and folded proteins and cross-validation following HX-based ensemble reweighting. **a**. Correlation between experimental (Exp.) and calculated (Calc.) PFs for disordered protein ensembles refined against NMR chemical shifts and SAXS data (Methods), using three representative forward models (BV2006, M9, and M14DH_scaled). **b**. Experimental and M14DH_scaled-predicted PFs for four representative disordered proteins. **c**. Concordance correlation coefficients (*ρ*_*c*_) of forward models for the NMR/SAXS-refined disordered protein ensembles. **d**. Helical propensities of (AAQAA)_3_ before (prior) and after HX reweighting (posterior) with three representative forward models, compared with experiment. **e**. Experimental and predicted PFs (left) and C^*′*^ secondary chemical shifts (right) of (AAQAA)_3_ before and after reweighting. RMSE values are shown for the prior, posterior 1, and posterior 2 ensembles. **f**. C*α* root-mean-square fluctuations (RMSF) of villin headpiece (HP36) before and after HX reweighting with three representative forward models. **g**. Experimental and predicted PFs (left) and C*α* secondary chemical shifts (right) of HP36 before and after reweighting. RMSE values are shown for the prior, posterior 1, and posterior 2 ensembles. **h**. Forward-model performance for HX data (*ρ*_*c*_ of predicted log_10_ PF) and cross-validation against orthogonal data for disordered (left) and folded (right) proteins. Positive and negative cross-validation scores indicate improved and worsened agreement, respectively, averaged across the dataset and all orthogonal data types (Methods).

### Cross-validation of HX-reweighted ensembles

We next used orthogonal NMR and SAXS observables to assess whether HX-driven reweighting improved the underlying structural ensembles, providing an independent measure of forward-model performance. For the helical peptide (AAQAA)_3_, reweighting against HX data with different forward models produced distinct changes in helical propensity (Figure 2d). Cross-validation against chemical shifts showed that M14DH_scaled yielded the best posterior ensemble, with improved agreement relative to the prior and a larger improvement than obtained with M9, whereas reweighting with BV2006 worsened agreement (Figure 2e). A similar pattern was observed for the folded villin headpiece HP36. Here, BV2006 increased backbone flexibility relative to the MD prior, whereas the explicit open-state models produced somewhat more rigid ensembles (Figure 2f). These changes were reflected in the chemical shifts, which wors-ened after BV2006 reweighting but improved with M9 and M14DH_scaled, though we note that the shifts are relatively small (Figure 2g).

Across the IDP dataset, explicit open-state models with electrostatic corrections also performed best after HX reweighting (Figure 2h, Supplementary Figure 5c), consistent with the analysis of independently NMR/SAXS-refined ensembles above. Overall agreement with orthogonal observables remained similar or improved after HX reweighting, indicating that better agreement with HX was not achieved through substantial deterioration of the broader ensemble properties. Chemical-shift agreement tended to improve for shorter systems such as (AAQAA)_3_, whereas for longer IDPs it was generally retained close to the prior level. In contrast, empirical forward models more frequently worsened agreement with SAXS and corresponding estimates of *R*_*g*_ (Supplementary Figure 5a).

For folded proteins, explicit open-state models likewise outperformed the empirical models after reweighting, although inclusion of the electrostatic correction had a smaller effect than for IDPs (Figure 2h, Supplementary Figure 5d), consistent with protection in folded proteins being dominated more strongly by structural burial and hydrogen bonding, and with folded proteins typically having fewer charged residues. More generally, explicit open-state models tended to maintain or modestly improve agreement with orthogonal data, whereas empirical models frequently degraded it, particularly for the more extensively reweighted posterior 2 ensembles (Figure 2h). This effect was driven largely by poorer agreement with NMR chemical shifts and amide order parameters after reweighting with BV2006 and BV2006ang, while the same observables were generally preserved or improved with explicit open-state models (Supplementary Figure 5b). Consistent with the HP36 example, empirical models tended to shift folded protein ensembles towards increased flexibility, whereas explicit open-state models produced slightly reduced flexibility on average, which correlated with improved agreement with NMR observables (Supplementary Figure 5d).

Together, these cross-validation analyses support the explicit open-state formulation as a more faithful basis for HX-guided ensemble refinement while avoiding overfitting. In particular, M11–M14DH_scaled perform consistently well across both disordered and folded proteins, improving agreement with HX while preserving or improving agreement with independent structural and dynamical observables. Thus, while empirical models such as BV2006 are powerful when the goal is to predict the free energy of opening from a ground-state ensemble where amides are predominantly in a closed state, explicit open models such as M14DH_scaled are more appropriate when the goal is to assess or refine ensembles that explicitly represent both closed and open states regardless of their relative populations.

### Refinement of protein folding free-energy landscapes

We next asked whether HX-guided reweighting could recover not only improved ground-state ensembles, but also the relative populations of distinct conformational states across complete folding landscapes. This analysis relies on the expectation that the most strongly protected residues exchange predominantly from globally unfolded conformations, such that their PFs contain information about the equilibrium between folded and unfolded states [45]. For 25 proteins with HX-NMR data, including 20 with experimental measurements of the global folding free energy, Δ*G*_fold_, we evaluated whether reweighting improved agreement with both HX PFs and the equilibrium between folded and unfolded states (Table 6, Methods). To generate ensembles spanning these landscapes, from highly disordered to natively folded conformations, we used BioEmu, a generative model that samples decorrelated protein conformations from an approximate room-temperature Boltzmann distribution [4]. This enabled broad folding landscapes to be sampled for proteins up to 87 residues in length (Figure 3a, Supplementary Figure 6a). In this dataset, all amides sampled an open state under every explicit model definition and were therefore included in the analyses for all models.

**Fig. 3.**
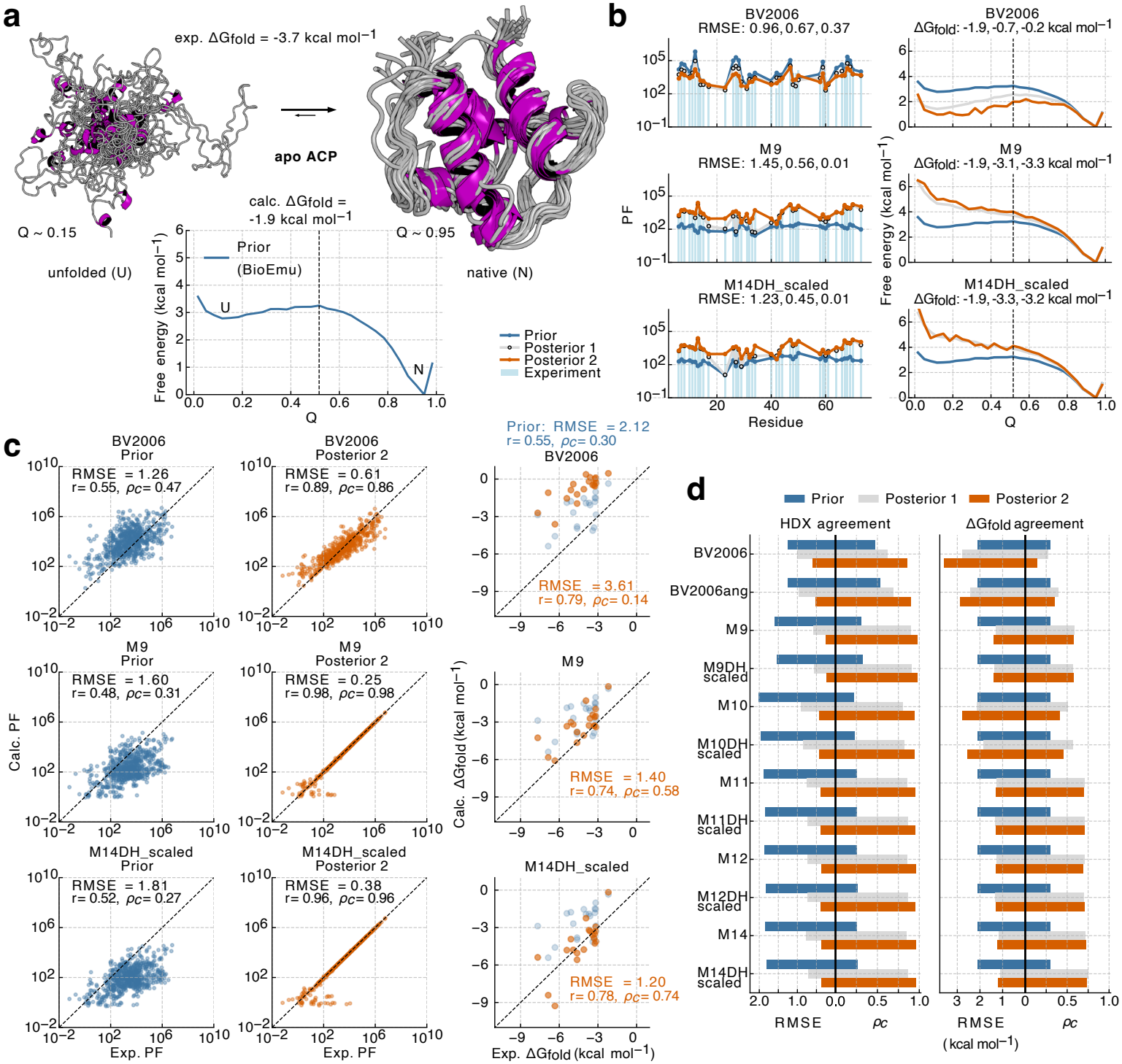
Evaluation of forward models on protein folding free-energy landscapes sampled with BioEmu. **a**. Representative unfolded and folded structures of apo acyl carrier protein (ACP), experimental and BioEmu-predicted folding free energies (Δ*G*_fold_), and the BioEmu-generated (prior) free-energy landscape as a function of the fraction of native contacts (*Q*). The vertical line indicates the cutoff separating unfolded and folded states, defined by the free-energy barrier between the two basins. **b**.Experimental and predicted PFs (left) and folding free-energy landscapes (right) of apo ACP before and after reweighting. RMSE and Δ*G*_fold_ values are shown for the prior, posterior 1, and posterior 2 ensembles. **c**. Correlation between experimental (Exp.) and calculated (Calc.) PFs across 707 amides from 25 proteins for three representative forward models (BV2006, M9, and M14DH_scaled), shown for the prior and posterior 2 ensembles. Agreement between experimental and calculated Δ*G*_fold_ values is shown for the subset of 20 proteins with measured folding stabilities (right). **d**. Forward-model performance for HDX data and cross-validation against experimental Δ*G*_fold_ values from panel c.

We then applied the same reweighting and cross-validation framework used for the ground-state ensembles. As an example, we show the results for apo acyl carrier protein (ACP), for which different forward models produced markedly different changes in the folding landscape (Figure 3a–b). Reweighting with BV2006 strongly increased the population of unfolded conformations, whereas the explicit open-state models shifted weight towards the folded state. This behaviour was representative of the broader dataset. BV2006 produced only limited improvement in agreement with HX-NMR PFs while substantially worsening agreement with experimental folding free energies, increasing the Δ*G*_fold_ RMSE from 2.1 to 3.6 kcal mol^*−*1^ after reweighting (Figure 3c–d). In contrast, the explicit open-state models, including M9, produced substantially closer agreement with HX-NMR while also yielding more realistic folded-state populations.

The best overall performance was obtained with models that include a packing-based descriptor, M11–M14DH_scaled. When we reweighted the folding landscapes generated by BioEmu using the HX data with M14DH_scaled, the resulting landscapes showed an improved agreement with folding free energies across a broad range of systems (RMSE dropped from 2.1 kcal mol^*−*1^ to 1.2 kcal mol^*−*1^, and *ρ*_*c*_increased from 0.30 to 0.74; Figure 3c). M9 also showed substantial improvement over the empirical models, but the improved performance of M11–M14DH_scaled suggests that the more stringent SASA and hydrogen-bond criteria together with directional packing provide additional information for identifying conformations that are genuinely protected from exchange. M10, which lacks the *ρ*_front_ packing criterion, tended to overstabilise folded conformations, resulting in a less favourable RMSE and only a modest improvement in *ρ*_*c*_ relative to the prior ensembles (Figure 3c–d, Supplementary Figure 6b–c). The electrostatic correction had only a minor effect in this dataset, consistent with structural burial and hydrogen bonding dominating protection in stably folded proteins.

We additionally tested whether representative forward models could recover native-state populations from deliberately poorer prior ensembles. For this purpose, we used long-timescale all-atom MD simulations of *α*3D, chignolin (CLN025), protein G (NuG2) and ubiquitin [46, 47], performed near their respective in silico melting temperatures such that folded and unfolded conformations were both substantially populated. These ensembles therefore provide a stringent test of whether HX-guided reweighting can shift a highly heterogeneous prior towards the experimentally relevant native basin (Supplementary Figure 6d–e). Consistent with the BioEmu analysis, reweighting with BV2006 produced little or no shift towards the folded state. M9 shifted the equilibrium more strongly towards native conformations, whereas M10– M14DH_scaled produced the largest enrichment of the folded basin. This provides an independent test that the newer explicit open-state models more accurately identify the conformations that should dominate the equilibrium ground-state ensemble.

These results show that the choice of HX forward model and structural definition of exchange competence directly affect the inferred populations of folded and unfolded conformations. While empirical models frequently shifted ensembles away from independently measured folding thermodynamics, explicit open-state models generally produced more accurate conformational populations. Importantly, however, the explicit models share the same functional relationship between protection and exchange-competent populations, yet produce different posterior ensembles. M9 already improves on the empirical models, while the newer M11–M14 variants perform better overall; however, M10 tends to over-stabilise native conformations, consistent with its less complete treatment of steric protection in the absence of the *ρ*_front_ packing term. Together with the analyses of disordered proteins and folded ground states, these results support the transferability of the M11–M14DH_scaled models across systems spanning a broad range of structural order and conformational heterogeneity. These results also highlight the importance of independent experimental validation for distinguishing among forward models that reproduce HX measurements but yield different conformational ensembles. In the remainder of this work, we thus focus on explicit open-state models.

### Millisecond HDX-MS captures conformational properties of disordered proteins

While HX-NMR provides highly accurate and residue-resolved measurements, it may be more difficult to apply to larger or scarcely available proteins. HDX-MS solves some of these issues, but provides coarser experimental information only at the level of individual peptides. We therefore extended the framework from residue-resolved HX-NMR protection factors to peptide-resolved HDX-MS measurements to combine informative structural priors with the less detailed measurements. For the explicit open-state models, residue-level PFs were converted into predicted peptide uptake curves by calculating the expected fraction of deuterated amides at each time point and summing over all exchangeable amides within a peptide, while assuming uniform back-exchange (Methods).

To test the accuracies of these calculations, we first focused on disordered peptides and proteins, for which protection is generally weak and even modest errors in predicted PFs translate directly into detectable deviations in peptide uptake kinetics. To this end, we used published HDX-MS measurements of model peptides, angiotensin and bradykinin [48]. Additionally, we collected millisecond HDX-MS datasets for A*β*42, *α*-synuclein and the 2N4R isoform of Tau (Methods). Sequence coverage was 100% for A*β*42 and *α*-synuclein and 89% for Tau (Supplementary Data). Despite their overall high exchange rates, different protein regions displayed clearly distinct uptake kinetics. For example, the C-terminal regions of A*β*42 and *α*-synuclein and the N-terminal region of Tau exchanged more slowly than other regions of the respective proteins (Figure 4a). In *α*-synuclein and Tau, these slower-exchanging regions are enriched in acidic residues (Supplementary Figure 7a), consistent with previous observations that local electrostatic environments can reduce hydroxide-catalysed amide exchange rates [40, 41].

**Fig. 4.**
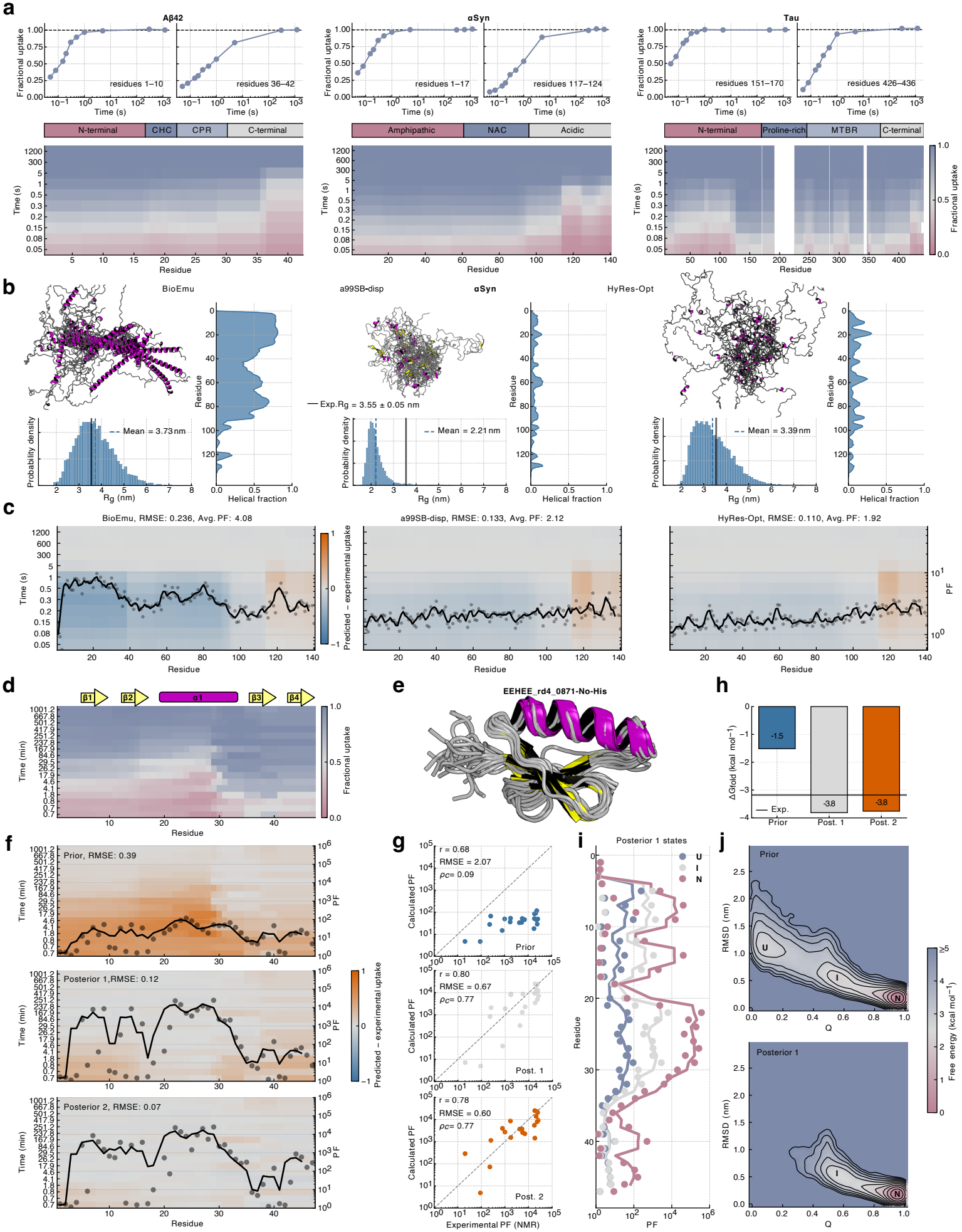
Integrative modelling with fast and conventional HDX-MS to benchmark and refine conformational ensembles of disordered and folded proteins. **a**. Millisecond HDX-MS data for three intrinsically disordered proteins, showing representative peptide uptake curves (top), domain organisation (middle), and uptake heatmaps across the protein sequence (bottom), where values at residues covered by multiple peptides are averaged across all overlapping peptides. **b**. Three *α*-synuclein ensembles generated with BioEmu, all-atom MD (a99SB-disp), and a hybrid resolution model (HyRes-Opt), showing representative structures, mean helical propensities, and radius of gyration (*R*_*g*_) distributions compared with experiment. **c**. Difference between predicted and experimental per-peptide deuterium uptake across time (heatmaps; left y-axis, overall RMSE indicated) and ensemble-averaged PFs (right y-axis with running average over three residues) for the three *α*-synuclein ensembles. Heatmaps show values across the protein sequence, with residues covered by multiple peptides averaged across all overlapping peptides. **d**. Conventional HDX-MS data for the designed protein EEHEE_rd4_0871 with native secondary structure indicated above. **e**. Representative BioEmu ensemble of the native EEHEE_rd4_0871 state. **f**. Difference between predicted and experimental per-peptide deuterium uptake across time (heatmaps; left y-axis) and ensemble-averaged PFs (right y-axis with running average over three residues) for the BioEmu-generated EEHEE_rd4_0871 ensemble before and after HDX-MS reweighting. **g**. Cross-validation against residue-resolved HX-NMR PFs before and after HDX-MS reweighting. **h**. Experimental and calculated folding free energies of EEHEE_rd4_0871 before and after reweighting. **i**. Calculated PFs for unfolded (U), intermediate (I), and native (N) conformational states after reweighting. **j**. Free-energy landscapes of EEHEE_rd4_0871 as a function of backbone RMSD from the native state and fraction of native contacts (*Q*) before and after reweighting.

To assess whether the relative performance of the explicit open-state models on these fast HDX-MS data was consistent with the HX-NMR analyses, we performed all-atom simulations for the model peptides, while we generated ensembles of the disordered proteins using the HyRes-Opt hybrid-resolution force field (Methods; Table 5). The IDP ensembles were refined against NMR chemical shifts and SAXS data and independently cross-validated against RDCs for all three proteins and PRE intensity ratios for *α*-synuclein and Tau (Supplementary Figures 4a and 7b). We then compared predicted and experimental peptide-level uptake values across the explicit open-state models M9–M14.

Because predicted HDX-MS kinetics of disordered systems are sensitive to the intrinsic exchange-rate reference, we performed the analysis using both the traditionally used poly-DL-alanine (PDLA) reference rates [49] and rates derived from the 3-Ala tripeptide, which were recently shown to improve H–D exchange rate predictions for disordered peptides [48]. Consistent with this work, we found that all forward models showed improved agreement with the HDX-MS data when using the 3-Ala reference rates (Supplementary Figure 7c). The relative ranking of forward models was also consistent with the HX-NMR results and with the model-peptide analysis above. M9 showed poorer agreement than M10–M14, consistent with its tendency to overpredict protection in short peptides (Supplementary Figure 3), whereas the more stringent SASA criteria used in M10–M14 more effectively distinguished transient solvent occlusion from genuine protection.

Electrostatic corrections further improved predictions for M10–M14, with M14DH_scaled reaching an average per-protein uptake RMSE of approximately 0.07 (Supplementary Figure 7c). Importantly, these structure-based predictions also outperformed a baseline model based solely on intrinsic exchange rates [48, 49], corresponding to a uniform *PF* = 1 for all amides. The same relative trends between forward models were retained when the conformational ensembles were reweighted directly against the HDX-MS data (Supplementary Figure 7d).

Together, these analyses identify two complementary contributions that are required for accurate prediction of fast HDX-MS kinetics in disordered systems. First, the stringent SASA, hydrogen-bonding and packing-density criteria of the M11–M14 models more accurately distinguish genuinely protected conformations from transient solvent occlusion; in contrast, M9 systematically overpredicts protection and consequently predicts uptake kinetics that are too slow (Supplementary Figure 7e–f). Second, explicit treatment of electrostatic effects captures longer-range modulation of hydroxide-catalysed exchange that is not fully represented by the local side-chain corrections contained in intrinsic-rate models. These results show that millisecond HDX-MS is sensitive to conformational and electrostatic features of IDP ensembles and can therefore provide quantitative constraints on their structural populations.

### Ranking and refining IDP conformational ensembles with HDX-MS constraints

Having established the transferability of the M11–M14DH_scaled forward models, we next asked whether millisecond HDX-MS could distinguish between alternative conformational ensembles of IDPs. IDP ensembles are notoriously sensitive to the underlying simulation force field [50] and generative AI model [11]. NMR and SAXS data have therefore been extensively used to parameterise force fields, benchmark generative models, validate structural ensembles and refine their conformational populations [11, 51–55]. We tested whether millisecond HDX-MS could provide analogous discriminatory power, while offering the potential for higher-throughput measurements with lower sample requirements [56].

As a proof of concept, we compared three distinct *α*-synuclein ensembles generated using BioEmu [4], long-timescale all-atom simulations with the a99SB-disp force field [57], and HyRes-Opt simulations (Table 5). These ensembles differed markedly in both secondary-structure content and global dimensions. BioEmu sampled extensive helical structure across the N-terminal ~90 residues, a99SB-disp predominantly sampled more compact conformations without persistent secondary structure, whereas HyRes-Opt generated a more extended and dynamic ensemble lacking stable secondary structure (Figure 4b). These differences were reflected in their agreement with independent NMR chemical shifts and SAXS data: BioEmu reproduced the chemical shifts poorly but showed good SAXS agreement, a99SB-disp agreed well with the chemical shifts but less well with SAXS, whereas HyRes-Opt agreed well with both observables (Supplementary Figure 8a).

The same ranking was reflected in the HDX-MS data using M14DH_scaled. HyRes-Opt showed the best overall agreement with experimental uptake, followed by a99SB-disp, whereas BioEmu showed substantial and systematic overprotection in the N-terminal region associated with its elevated helical propensity (Figure 4c). Residual disagreement at the *α*-synuclein C-terminus for the HyRes-Opt ensemble depended on the scaling of the atomic charges used in the electrostatic correction (Supplementary Figure 8b), suggesting that electrostatic effects not fully captured by either the ensemble or forward model contribute to the remaining discrepancy. We further reweighted the BioEmu and a99SB-disp ensembles directly against the HDX-MS data. Reweighting improved agreement with chemical shifts for BioEmu and with SAXS for a99SB-disp, accompanied by a reduction in excessive helical structure in BioEmu and overly compact conformations in a99SB-disp (Supplementary Figure 8c–d). Thus, HDX-MS was able not only to distinguish between alternative *α*-synuclein ensembles, but also to refine structural features that were independently supported by orthogonal measurements.

We observed similar behaviour for the other IDPs. For A*β*42, we compared ensembles generated with the atomistic a99SB*-ILDN force field [58, 59], HyRes-GPU [60] and HyRes-Opt (Table 5). Agreement with HDX-MS using M14DH_scaled tracked agreement with chemical shifts and the corresponding secondary-structure propensities, with the HyRes-Opt ensemble, which sampled relatively extended conformations lacking persistent secondary structure, providing the best overall agreement (Supplementary Figure 8e–f). For Tau, we compared an ensemble generated previously using the IDP-o model [11] with HyRes-GPU and HyRes-Opt ensembles. Again, agreement with HDX-MS broadly followed the independent NMR and SAXS assessments. HyRes-Opt sampled a broad ensemble of extended conformations with little persistent secondary structure and showed good agreement with chemical shifts, SAXS and HDX-MS. In contrast, the more compact HyRes-GPU ensemble showed poorer agreement with both SAXS and HDX-MS, while IDP-o showed poorer agreement with chemical shifts and HDX-MS, consistent with its higher helical propensity across the protein (Supplementary Figure 8g–h).

Together, these results show that millisecond HDX-MS, when interpreted using an appropriate forward model, can benchmark, rank and refine conformational ensembles of IDPs in a manner that is consistent with independent NMR and SAXS measurements. Across the ensembles examined here, agreement with HDX-MS correlated particularly strongly with agreement with NMR chemical shifts, consistent with hydrogen bonding and local secondary structure being major determinants of amide protection. Remarkably, however, differences in global compaction also produced clear changes in HDX-MS agreement, indicating that peptide uptake is sensitive not only to local structure but also to longer-range conformational organisation of IDP ensembles. Crucially, the inclusion of both structural and electrostatic terms in the forward model provides an opportunity to deconvolute their respective contributions to protection, which would not be apparent from analysis of the experimental HDX-MS, or HX-NMR data [44], alone.

### Integrative modelling with HDX-MS captures a folding intermediate

We next applied HDX-MS-guided reweighting to a folded domain, the designed protein EEHEE_rd4_0871 [14], using a previously published HDX-MS dataset [61]. Experimentally, the central helical region showed the slowest deuterium uptake, whereas the two C-terminal *β*-strands exchanged more rapidly than the two N-terminal strands (Figure 4d–e). The initial BioEmu ensemble predicted uptake that was too rapid, reflecting insufficient stability of the N-terminal strands and central helix. Reweighting substantially improved agreement with the HDX-MS data by increasing protection in these regions, while the C-terminal strands remained the least protected part of the protein (Figure 4f, Supplementary Figure 9a).

We selected EEHEE_rd4_0871 to assess our protocol because residue-resolved NMR protection factors had been measured independently [14], thus enabling us to assess how well reweighting against HDX-MS data could recover residue-specific information. We find that the agreement with these NMR PFs improved markedly after HDX-MS reweighting, even for the minimally perturbed posterior 1 ensemble, with *ρ*_*c*_ increasing from 0.09 to 0.77; more extensive reweighting in posterior 2 produced no further improvement (Figure 4g). The predicted global stability likewise moved into substantially better agreement with an independent limited-proteolysis measurement [62], again without further improvement from posterior 1 to posterior 2 (Figure 4h). These results show that reweighting against peptide-level HDX-MS data leads to an accurate description both at the local level (residue-specific PFs) and global level (folding free-energies). Because the results were similar for the posterior 1 and 2 ensembles, we focused subsequent structural analysis on the posterior 1 ensemble.

Inspection of the original BioEmu free-energy landscape revealed a distinct folding intermediate, corresponding to a local minimum in addition to the unfolded and folded states. This intermediate persisted after reweighting, while the population of the unfolded state decreased (Figure 4i–j). Consistent with the experimentally faster deuterium uptake in the C-terminal region, the intermediate retained protection in the first two *β*-strands and central helix, whereas the C-terminal strands remained essentially unprotected and indistinguishable from the unfolded state (Figure 4i). This pattern was also consistent with the independently measured NMR PFs, which showed the weakest protection in the C-terminal region [14]. Reweighting the same ensemble directly against the NMR PFs produced a highly similar free-energy landscape, including an intermediate populated at approximately 5% (Supplementary Figure 9a).

Structurally, conformations belonging to this intermediate showed an almost complete loss of secondary structure in the C-terminal region, while the remainder of the protein remained largely native-like (Figure 5a, Supplementary Figure 9a). Together, these results provide a proof of concept that computational ensembles and HDX-MS can be integrated not only to improve ensemble populations, but also to structurally resolve and quantify partially folded intermediates within heterogeneous conformational landscapes. While the accuracy of the results depend on whether the prior ensemble captures the relevant conformational states, the integration with the experimental data enables us to combine the power of generative models to sample relevant states with experiments to recover an improved thermodynamic landscape.

**Fig. 5.**
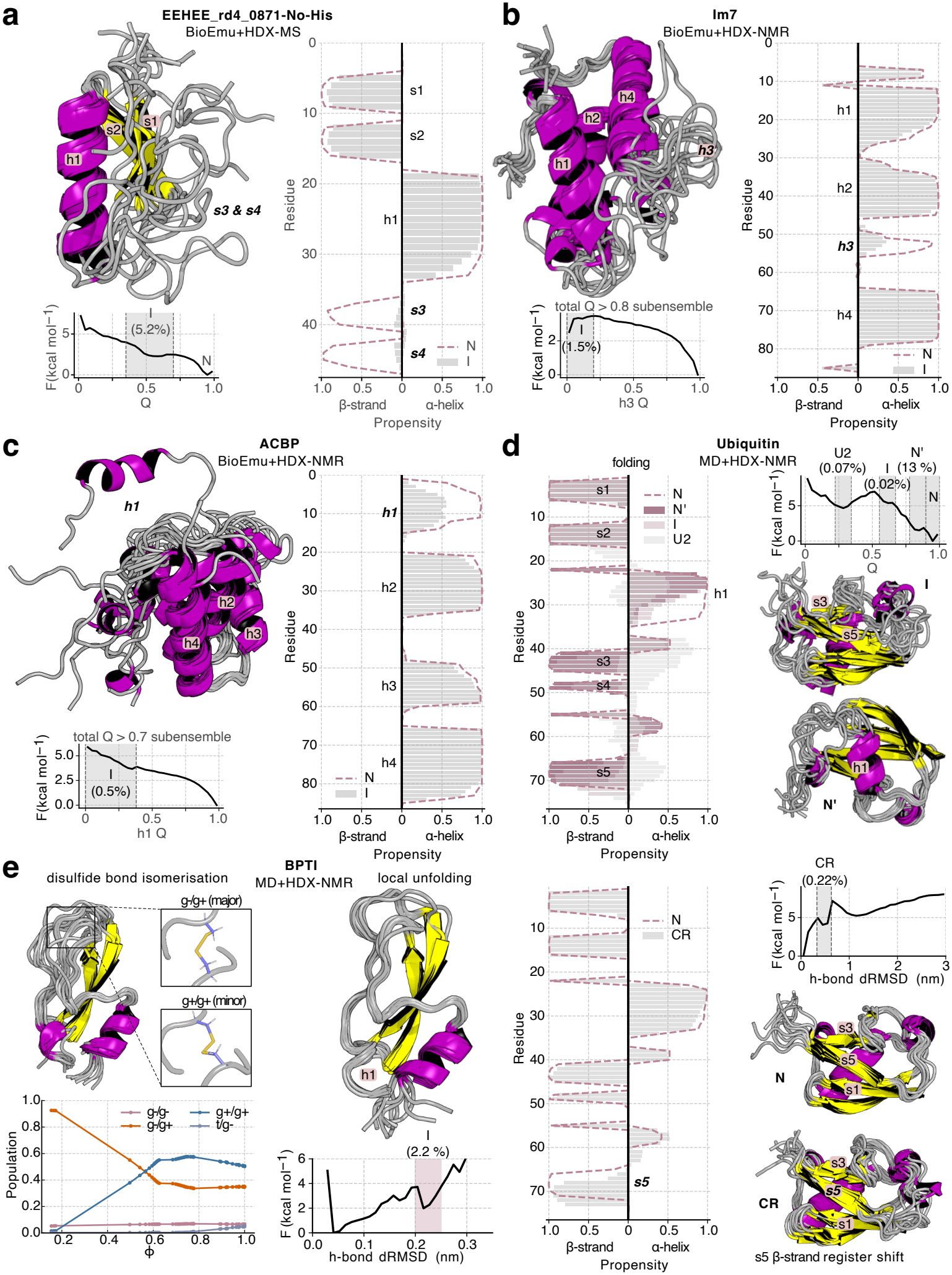
Folding intermediates and excited conformational states identified from HDX-reweighted ensembles. **a**. Ensemble of the EEHEE_rd4_0871 intermediate (I) state after HDX-MS reweighting, its secondary structure propensities compared to the native (N) state and the free-energy landscape along the fraction of native contacts (*Q*). **b**. Ensemble of the Im7 I state after HDX-NMR reweighting, its secondary structure propensities compared to the N state and free-energy profile along helix 3 (h3) native contacts within the folded (*Q >* 0.8) subensemble. **c**. Ensemble of the ACBP I state after HDX-NMR reweighting, its secondary structure propensities compared to the N state and free-energy profile along helix 1 (h1) native contacts within the folded (*Q >* 0.7) subensemble. **d**. Ubiquitin MD ensemble reweighted with HDX-NMR data, showing secondary structure propensities and representative structures of conformational states along the folding free-energy landscape using *Q* (top), and a non-native excited state characterised by a C-terminal *β*-strand s5 register shift (CR, C-terminally retracted state) identified along a one-dimensional free-energy profile of s5 hydrogen bond distance RMSD (dRMSD) relative to the N state. **e**. Representative MD structures of BPTI in the experimentally identified major state (Cys14-Cys38 disulfide g+/g+ rotamer) reweighted with HDX-NMR data. Disulfide bond rotamer population changes during reweighting are shown (as a function of *ϕ*, bottom left). A locally unfolded (helix 1, h1) excited state characterised by an increased hydrogen bond dRMSD relative to the N state is also identified with representative structures (top, right).

### Intermediates and excited states across proteins

To further test whether HX-reweighting can identify folding intermediates and low-population excited states, we analysed additional proteins for which such states have been characterised independently. For Immunity protein 7 (Im7), reweighting improved agreement with independent measurements of global protein stability [63] and revealed a folding intermediate in which helices 1, 2 and 4 remain folded and form a hydrophobic core, whereas helix 3 is detached and partially unfolded (Figure 5b, Supplementary Figure 9b). This local free-energy minimum was already predicted in the prior BioEmu ensemble and persisted after reweighting, consistent with the lower NMR PFs observed for helix 3 relative to the remaining helical regions [63]. A structurally similar intermediate had previously been proposed from *ϕ*-value analysis [64], and its population of approximately 1.5% is in good agreement with the experimental free-energy difference between the native and intermediate states [63, 65] (Supplementary Figure 9b).

For Acyl-CoA-Binding Protein (ACBP), NMR PFs are reduced in the N-terminal helix relative to the other helices [66]. BioEmu sampled conformations in which helix 1 is unfolded while the remainder of the protein remains largely native-like, and reweighting against the HX-NMR protection factors sharpened this region into a distinct local minimum corresponding to a partially detached and unfolded helix 1 with a population of approximately 0.5% (Figure 5c, Supplementary Figure 9c). This minimum was most pronounced in the more extensively reweighted posterior 2 ensemble (*ϕ* ~ 0.5), which also showed improved agreement with the global stability of ACBP [66] (Supplementary Figure 9c). The N-terminal helical region of ACBP has been shown to be stabilised upon ligand binding [66], consistent with a population shift away from the partially unfolded state detected here and towards the native conformation. A related intermediate has also been suggested from temperature-dependent NMR chemical shifts, which implicated perturbations of helices 1 and 3 [67]. Our HX-reweighted ensemble therefore captures some features of this previously proposed intermediate, although we do not resolve a second distinct minimum involving helix 3. The ensemble nevertheless samples conformations in which helix 3 is perturbed while the remainder of the protein remains largely native-like, suggesting that an additional low-population state may be present but is not sufficiently stabilised by the HX restraints to emerge as a separate free-energy minimum (Supplementary Figure 9c).

We next analysed ubiquitin using an extensive MD dataset that samples reversible folding transitions together with fully and partially unfolded conformations [47]. Because these simulations were performed at elevated temperature, the prior ensemble required more substantial reweighting than the BioEmu ensembles discussed above. Nevertheless, agreement with the independently measured global folding stability improved markedly, from 0.5 kcal mol^*−*1^ in the prior to −6.1 kcal mol^*−*1^ in posterior 2, compared with the experimental value of −6.8 kcal mol^*−*1^ [34] (Supplementary Figure 9d). The reweighted ensemble retained several metastable states also identified in the original MD study (Figure 5d). These included the U2 state, characterised by a folded N-terminal hairpin resembling residual structure observed by NMR under denaturing conditions [68]; a native-like intermediate with an established core but incompletely formed secondary structure involving helix h1 and strands s3–s5; and an N’-like state in which the C-terminal residues of helix h1 are locally unfolded, consistent with high-pressure NMR observations [69]. These structural features agree with the HX-NMR protection pattern, in which strands s1–s2 and the N-terminal portion of h1 are most strongly protected, whereas the C-terminal portion of h1 and strands s3–s5 show lower protection and greater conformational dynamics [34, 70] (Supplementary Figure 9d).

The MD ensemble also sampled a non-native excited state implicated in Ser65 phosphorylation by PINK1, characterised by a register shift of the C-terminal strand s5. This state remained populated at approximately 0.1–0.2% after HX reweighting (Figure 5d) and has been observed independently by pressure-jump NMR [71] and chemical exchange saturation transfer (CEST) NMR, with an estimated upper-bound population of approximately 0.7% at room temperature [72]. The populations of these non-native ubiquitin states were generally consistent between posterior 1 and posterior 2. The main exception was the N’ state, for which improved agreement with the h1 PFs in posterior 2 was accompanied by a reduced population of approximately 3.4% (Supplementary Figure 9d).

Finally, we reweighted a one-millisecond MD trajectory of natively folded BPTI [73] against HX-NMR PFs previously compiled by Persson and Halle [28]. A key structural feature of BPTI is the disulfide bond between Cys14 and Cys38. NMR relaxation-dispersion experiments have shown that this bond predominantly adopts a right-handed conformation, with Cys14 and Cys38 in gauche− /gauche+ (g− /g+) conformations, whereas a left-handed g+/g+ conformation forms an excited state populated at approximately 1.4% near room temperature [74]. In contrast, the MD force field overpopulated the g+/g+ state, which accounted for approximately 50% of the prior ensemble. The different disulfide conformers exhibited subtle differences in predicted HX protection, primarily around the disulfide residues and the C-terminal helix h2 (Supplementary Figure 9e). Remarkably, despite these protection differences being much smaller than those associated with the partially or globally unfolded states analysed above, HX reweighting shifted the ensemble towards the experimentally dominant g− /g+ conformer, with the g+/g+ population falling below that of the native conformer once *ϕ* decreased below approximately 0.55 (Figure 5e). The trajectory also sampled a metastable state involving local unfolding of the N-terminal helix h1. This state was retained after reweighting at a population of about 2.2% and was characterised by disruption of hydrogen bonds within h1 while the helix remained packed against the protein core (Figure 5e, Supplementary Figure 9e). Together, the BPTI analysis shows that HX restraints can resolve and reweight comparatively subtle native-state heterogeneity, in addition to the more pronounced folding intermediates and partially unfolded states identified above.

## Discussion

Here we establish a framework for quantitatively integrating HX measurements with conformational ensembles across disordered, folded and partially unfolded proteins. Across the systems examined, explicit open-state models provided a more transferable description of HX than empirical protection models, in particular for integrative modelling. Importantly, HX-driven reweighting with these models generally preserved or improved agreement with independent NMR, SAXS and folding-stability measurements, indicating that improved agreement with HX reflected more realistic conformational populations rather than simply a better fit to the restrained data.

The best-performing models used simple and physically motivated structural descriptors—hydrogen bonding, solvent accessibility and directional steric packing— together with electrostatic modulation of exchange. Their performance was relatively robust to modest changes in SASA thresholds, the precise definition of the packing descriptor and charge scaling: M11, M12 and M14 with electrostatic corrections performed similarly across the main benchmarks. M14DH_scaled was therefore used primarily as a representative (default) model rather than as an optimal parametrisation. This robustness suggests that the principal requirement is an appropriate physical definition of exchange competence rather than fine-tuning of individual structural cut-offs. More complex approaches, including machine-learning models, may ultimately improve forward-model accuracy as larger HX datasets become available [14]. However, the scarcity of experimentally determined ground-truth ensembles [5] and the strong correlations between possible structural descriptors create a substantial risk of overfitting, making validation against observables not used to construct the model particularly important.

Importantly, our results demonstrate that the functional form of a forward model is not sufficient to determine its suitability for ensemble refinement. Explicit open-state models with the same underlying relationship between protection and opening probability differed in their predictions of HX in disordered proteins and peptides, and produced different conformational populations when reweighted against the same experimental data. These differences were particularly evident in the independent folding-stability validation, where models yielding good agreement with HX did not necessarily recover the correct folded-state populations. Together, these findings highlight the importance of both the physical definition of exchange competence and independent experimental validation when developing HX forward models.

Our results also extend the quantitative use of HDX-MS beyond the conventional interpretation of relative protection patterns [75]. Millisecond HDX-MS distinguished alternative IDP ensembles and, upon reweighting, shifted structural properties towards those independently supported by NMR and SAXS. For folded proteins, HX provided strong sensitivity to partially unfolded and low-population conformations, including intermediates in EEHEE_rd4_0871, Im7 and ACBP, and native-state heterogeneity in ubiquitin and BPTI. This sensitivity arises because strongly protected amides may exchange predominantly through rare open conformations, making HX particularly informative about states that contribute only weakly to many conventional ensemble averages [12, 13]. Quantitative integration of HDX-MS with computational ensembles therefore complements established approaches for ensemble refinement [30–32] and previous efforts to reweight simulated ensembles against peptide-level HDX-MS measurements [24–26].

Two limitations remain important. First, reweighting can only modify populations of conformations already represented in the prior ensemble [76]; relevant opening events or excited states that are not sampled cannot be recovered. This will become increasingly restrictive for large proteins and complexes, where comprehensive sampling remains difficult [4]. Second, HX measurements do not uniquely determine a microscopic conformational ensemble. As with other ensemble-averaged measurements, many distributions may reproduce the same experimental observables [76]. HX is therefore likely to be most powerful in integrative applications alongside complementary measurements such as NMR, SAXS, single-molecule experiments [77] and single-particle cryo-EM [78], which constrain different aspects of conformational heterogeneity.

More broadly, the states identified here highlight how much structural heterogeneity remains hidden behind conventional single-structure descriptions and predictions of proteins [1, 14]. Transient structure in IDPs [55, 79], folding intermediates [7, 18, 19, 71] and low-population native-state conformations [9, 74] appear throughout the systems examined and may be functionally important [7, 9, 69, 72]. As molecular simulations and generative models increasingly enable broader and more accurate exploration of conformational space, scalable experimental measurements capable of evaluating these ensembles will become correspondingly important. The combination of HDX-MS with integrative ensemble refinement and accurate forward models such as those developed here provides a route towards uncovering this dynamic structural proteome and bench-marking and improving computational models of protein conformational ensembles at scale.

## Methods

### Literature hydrogen exchange NMR (HX-NMR) data

Datasets of proteins with hydrogen exchange NMR data (HX-NMR) were curated from the literature to assess different forward models across a large range of proteins, summarised in Table 1. Literature datasets were selected when residue-specific backbone amide hydrogen-exchange rates or protection factors were reported under sufficiently well-defined experimental conditions to permit calculation of intrinsic exchange rates and direct comparison with structural ensembles. We prioritised proteins for which the sequence and structural state were unambiguous, no ligands were present in the experiments (except structural ions) and for which complementary experimental data reporting on the ground-state structure or dynamics were available, while including both folded and intrinsically disordered proteins to ensure a broad range of protection.

Datasets were restricted, where possible, to measurements for which EX2 exchange had been experimentally demonstrated. Evidence for EX2 included an approximately linear dependence of the observed exchange rate on hydroxide concentration, with unit slope in the base-catalysed regime, or agreement with independently measured folding and closing kinetics showing that conformational closing was substantially faster than intrinsic chemical exchange. In several studies, pH-, temperature- or denaturant-dependent measurements provided additional evidence distinguishing EX2 exchange from EX1 or mixed exchange behaviour. For some datasets, assignment to the EX2 regime was supported by HDX-MS measurements of the same protein under comparable conditions, which showed unimodal isotopic envelopes and no detectable signatures of cooperative EX1 exchange. Where such tests were not reported, we selected measurements performed under native or near-native solution conditions, generally near neutral pH and approximately 300 K, and excluded conditions associated with substantial denaturation or reported EX1 behaviour. Under these conditions, the intrinsic chemical exchange rate is generally slower than conformational reclosing in stably folded proteins. An amide that becomes transiently exposed is therefore more likely to return to the protected state than to exchange during a single opening event, which is the defining kinetic condition for EX2 exchange. These conditions favour EX2 behaviour but do not by themselves demonstrate that every residue exchanges exclusively in the EX2 regime.

Intrinsic exchange rates were calculated using the poly-DL-alanine (PDLA) reference rates reported by the Englander lab (Nguyen et al. [49]). When experimental exchange rates were not reported directly, the rates were first reconstructed from the published protection factors using the reference-rate model employed in the original study, for example, that of Bai et al. [16, 80]. Protection factors were then recalculated consistently for all datasets using the Nguyen et al. reference rates according to

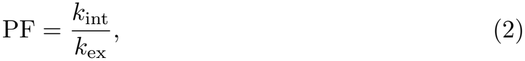

where *k*_int_ is the intrinsic exchange rate and *k*_ex_ is the experimentally measured exchange rate. For ensemble-reweighting calculations, all protection factors were assigned a relative uncertainty of 50% to account for uncertainty in the calculated intrinsic exchange rates and the heterogeneous and often incomplete reporting of experimental uncertainties across the literature datasets.

In addition, experimental data providing independent information on the ground-state structure and dynamics of each protein were collected from the literature for cross-validation analyses (Table 1). These orthogonal datasets included a range of solution NMR measurements as well as small-angle X-ray scattering (SAXS) data. For proteins whose sampled ensembles extended beyond the ground state and were used to analyse folding free-energy landscapes, experimentally determined folding stabilities were also collected (Table 6).

### Proteins and reagents

Human *α*-synuclein was purchased from Abcam as a 143 *µ*M stock, human Tau-441 (2N4R) from ACROBiosystems as a lyophilised powder, and human A*β*42 from GenScript as a 1 mg lyophilised aliquot. Sodium phosphate, NaCl, tris(2-carboxyethyl)phosphine (TCEP), guanidine hydrochloride (GdnHCl), and D_2_O (99.9 atom % D) were purchased from Sigma-Aldrich; formic acid (LC–MS grade) and acetonitrile (Optima LC–MS grade) from Fisher Scientific.

### Hydrogen-deuterium exchange mass spectrometry (HDX-MS)

We performed HDX-MS experiments on *α*-synuclein, A*β*42 and Tau. A*β*42 and Tau were reconstituted in equilibration buffer (20 mM sodium phosphate, 150 mM NaCl, 1 mM TCEP, pH 7.4) to 443 *µ*M (2.0 mg/mL) and 20.9 *µ*M (1.0 mg/mL), respectively. Immediately before HDX labelling, all proteins were diluted in equilibration buffer to their working concentrations (*α*-synuclein: 5.6 *µ*M; Tau: 5.1 *µ*M; A*β*42: 10 *µ*M). Deuterium labelling was performed using a Fast-HDX stopped-flow system (Applied Photophysics) coupled to two Waters Acquity I-class UPLC pumps and a Waters Cyclic IMS mass spectrometer. Labelling was initiated by 1:15 dilution into equilibration buffer prepared in 99.9% D_2_O (pD 7.4) at 20 °C, yielding a final D_2_O concentration of 93.3%. The following labelling time points were measured: 50, 80, 150, 200, 300, and 500 ms; 1 and 5 s; and 5 and 20 min for Tau and A*β*42, with an additional 10 min point for *α*-synuclein. At each time point, exchange was quenched by rapid addition of ice-cold quench buffer (2 M GdnHCl, 0.8% formic acid, pH 2.0) at 3 °C. Mixing, labelling, quenching, and injection were fully automated.

Quenched samples were delivered to an online digestion column maintained at 15 °C. *α*-synuclein was digested with an immobilised pepsin column (AffiPro). A*β*42 and Tau were digested with an immobilised nepenthesin-2 column (AffiPro). The digestion column was flushed twice between injections with 1.5 M GdnHCl, 0.8% formic acid in 96:4 H2O/acetonitrile to eliminate carryover. Peptides were trapped and desalted on an Acquity UPLC BEH C18 VanGuard pre-column (130 Å, 1.7 *µ*M, 2.1 × 5 mm; Waters) at 150 *µ*L/min for 4 min, then separated on an Acquity UPLC HSS T3 column (100 Å, 1.8 *µ*M, 1.0 × 50 mm; Waters) using a 5–35% acetonitrile gradient (0.1% formic acid) over 6 min at 100 *µ*L/min. The trap and analytical columns were maintained at 0.0 ± 0.1 °C throughout. Mass spectra were acquired on a Waters Cyclic IMS instrument in HDMSE mode, scanning 50–2000 m/z with ion mobility separation. We used the following source parameters: capillary voltage 2.80 kV, sampling cone 30 V, source temperature 80 °C, desolvation temperature 175 °C, trap and transfer collision energies 6 V and 4 V, respectively. Lock mass correction used glu-fibrinopeptide (200 fmol/*µ*L; Sigma-Aldrich). The instrument was calibrated with Major Mix solution (Waters) prior to data collection.

Peptides were identified using PLGS 3.0.1 (Waters) from undeuterated control acquisitions and imported into DynamX 3.0 (Waters) with filtering criteria of at least two consecutive fragment ions and a minimum of 0.25 fragment ions per amino acid. All spectra were manually inspected for correct isotopic centroid assignment. Sequence coverage, peptide count, and redundancy for each protein are reported in Supplementary Data. Centroid masses from different charge states were first averaged within each independent measurement. Undeuterated measurements were pooled for each peptide to obtain a reference mass, which was subtracted from the deuterated centroid mass to calculate absolute deuterium uptake. Absolute uptake values were subsequently averaged across independent measurements at each time point. The maximum deuterium uptake for our disordered protein data, maxD, was estimated separately for each peptide by fitting the mean uptake curve to a stretched exponential,

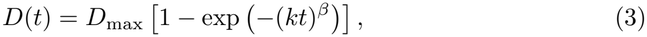

where *D*_max_ is the fitted plateau, *k* is an apparent exchange rate, and *β* is the stretching exponent. Fractional uptake was then calculated as *D*(*t*)*/D*_max_. A common experimental uncertainty was estimated by pooling the variance of replicate-level fractional uptake values across all peptide–time-point groups per protein,

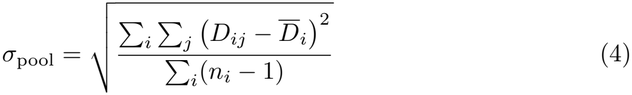

where *i* denotes a peptide–time-point group, *j* denotes an independent replicate, and *n*_*i*_ is the number of replicates in group *i*. This pooled standard deviation was assigned to all fractional uptake measurements. All HDX-MS datasets analysed in this study exhibited isotopic-envelope patterns consistent with exchange occurring predominantly in the EX2 regime.

Between one and four independent acquisitions were performed per time point for each protein. Replicate details, experimental parameters, and all processed values are provided in Supplementary Data, following community recommendations. (Masson et al., 2019). The HDX-MS data have been deposited to the ProteomeX-change Consortium via the PRIDE partner repository [81] with the dataset identifier PXD086174.

Additional datasets were collected from the literature. Table 2. All HDX-MS datasets analysed in this study exhibited isotopic-envelope patterns consistent with exchange occurring predominantly in the EX2 regime.

**Table 2:**
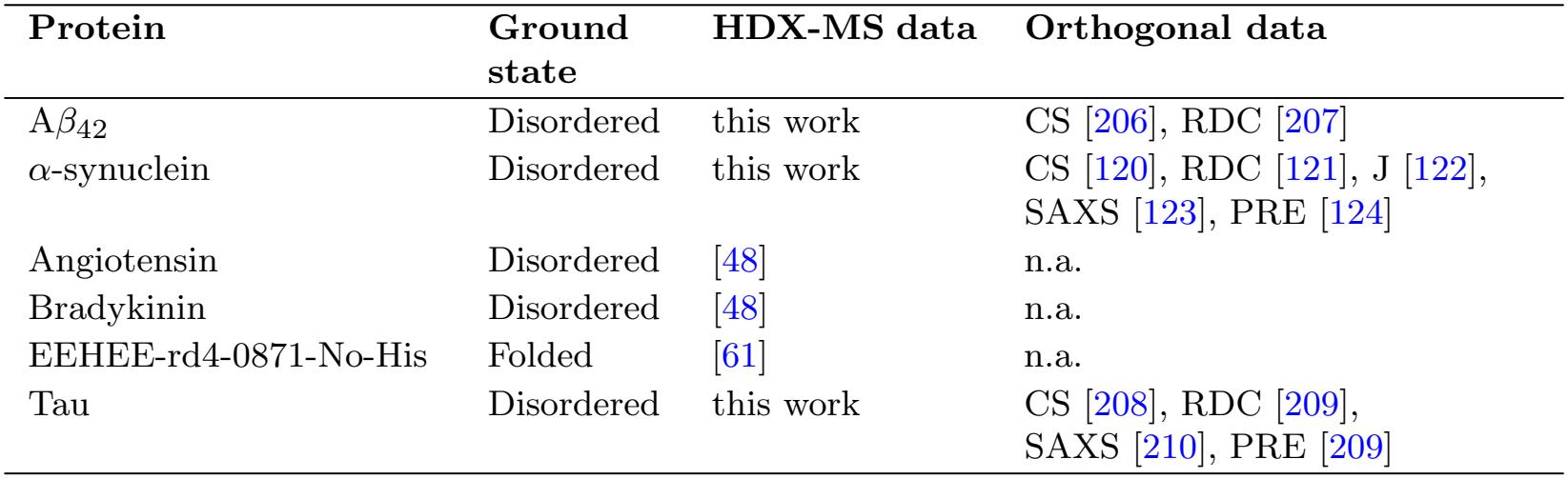
HDX-MS datasets and corresponding orthogonal experimental data reporting on structural and dynamic properties of the ground state. CS, backbone chemical shifts; RDC, amide residual dipolar couplings; J, backbone ^3^J-couplings; PRE, paramagnetic relaxation enhancement; SAXS, small-angle X-ray scattering; NOE, nuclear Overhauser effect distances; S^2^, amide order parameters; n.a., not applicable.

| **Protein** | **Ground state** | **HDX-MS data** | **Orthogonal data** |
|---|---|---|---|
| Aβ<sub>42</sub> | Disordered | this work | CS [206], RDC [207] |
| α-synuclein | Disordered | this work | CS [120], RDC [121], J [122],<br>SAXS [123], PRE [124] |
| Angiotensin | Disordered | [48] | n.a. |
| Bradykinin | Disordered | [48] | n.a. |
| EEHEE-rd4-0871-No-His | Folded | [61] | n.a. |
| Tau | Disordered | this work | CS [208], RDC [209],<br>SAXS [210], PRE [209] |

### All-atom molecular dynamics simulations

We used GROMACS versions 2023.1, 2023.4 and 2024.4 [82] for all-atom MD simulations. All proteins and peptides were placed in the centre of a dodecahedral-shaped box 1 nm from the box edges. Input structures, force fields, the number of solvent molecules and additional ions for each system are listed in Table 3. All systems were energy-minimised with the steepest descent algorithm, followed by a 1 ns equilibration in the NVT ensemble and 1 ns equilibration in the NPT ensemble. All dynamics simulations were performed using the leap-frog integrator, and LINCS [83] was used to constrain all covalent bonds involving hydrogen. Short-range van der Waals and electrostatic interactions were calculated using cutoffs of 0.9 and 1.0 nm, respectively, and particle mesh Ewald (PME) [84] was used for long-range electrostatic interactions with cubic interpolation and a Fourier spacing of 0.125 nm. The temperature was controlled with the velocity rescale method and a coupling time constant of 0.1 ps [85]. The pressure was kept at 1 bar using the Berendsen barostat [86], a compressibility of 4.5×10^*−*5^ bar^*−*1^ and a coupling constant of 2 ps during equilibration. The pressure coupling was subsequently switched to the Parrinello-Rahman barostat [87]. For most production simulations (Table 3), we employed a timestep of 4 fs using hydrogen mass repartitioning (HMR) [88].

We also performed all-atom molecular dynamics simulations of capped tripeptides, Ace–AXA–NMe, for all amino-acid types except proline. These simulations were used to characterise the solvent accessibility of the central amide associated with residue X in minimal model peptides resembling those used experimentally to determine intrinsic exchange rates. These simulations were performed at 300 K and 1 bar in the NPT ensemble with identical equilibration procedures as described above. The a99SB-disp force field [52] was used. A single 1 *µ*s simulation was performed for each tripeptide with normal atomic masses and a 2 fs timestep.

In addition to these in-house simulations described above and summarised in Table 3, we also used all-atom MD datasets from previously published works summarised in Table 4.

**Table 3:**
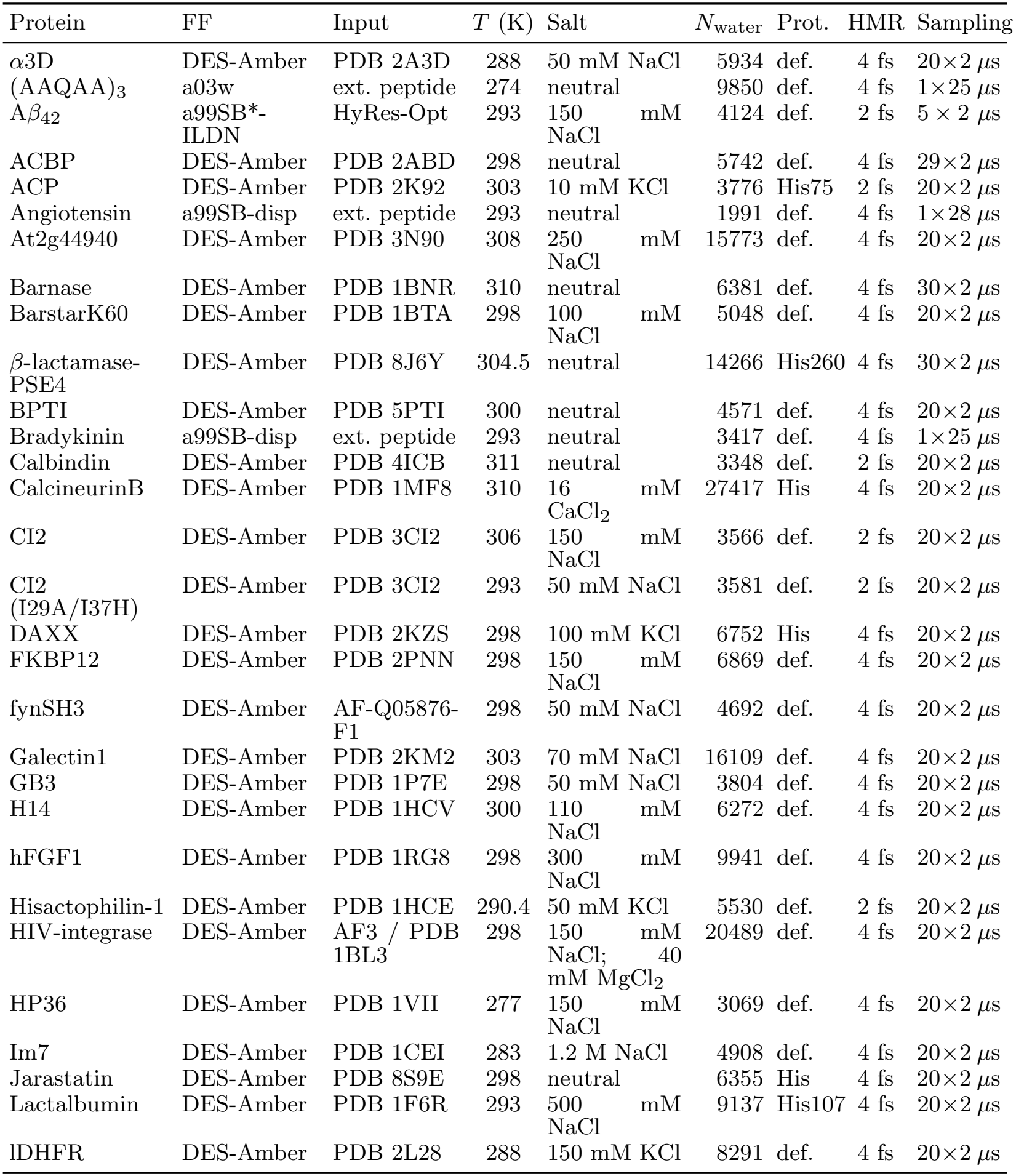
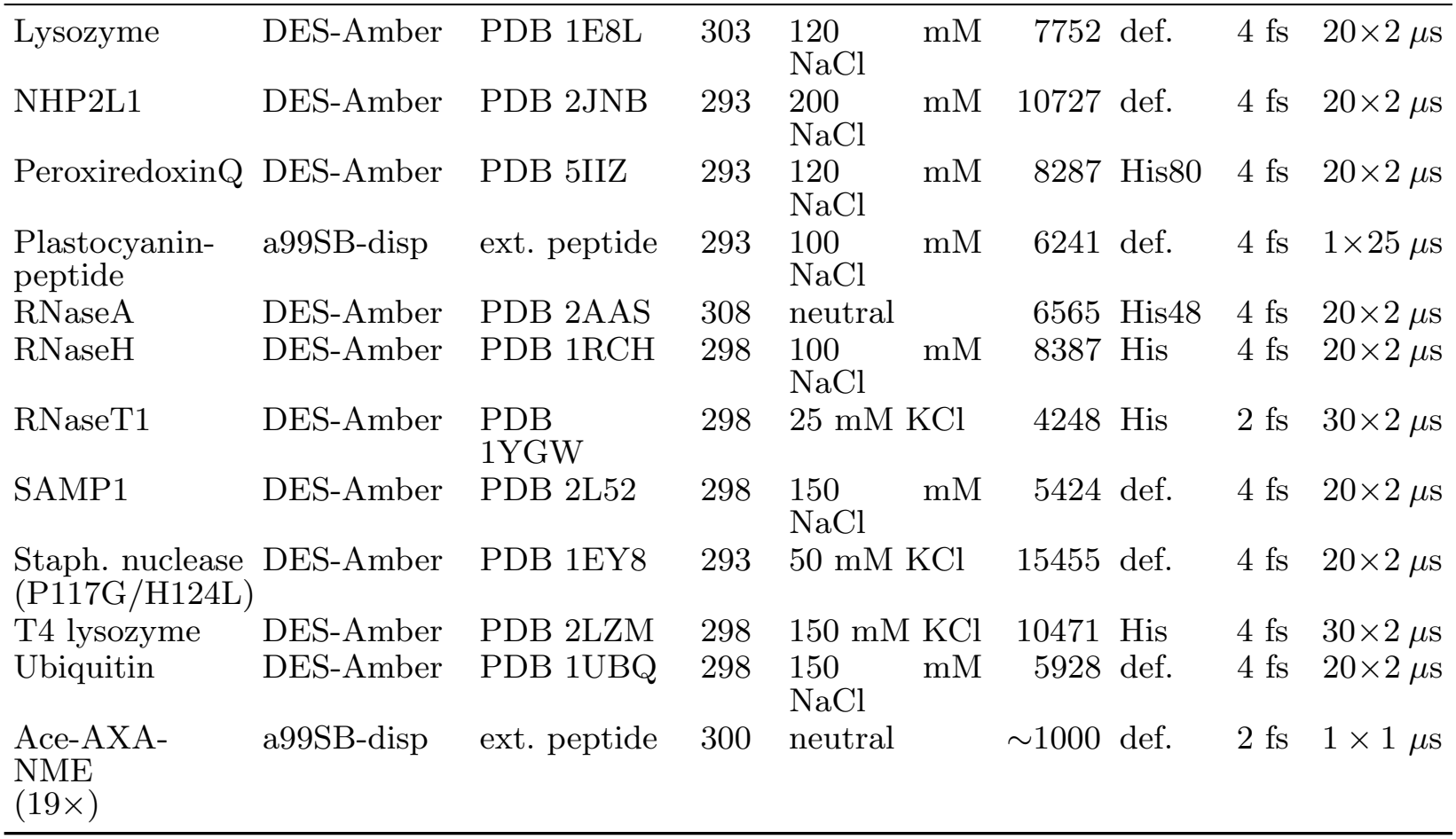
All-atom simulation systems and conditions. FF, force field; Prot., protonation; HMR, hydrogen mass repartitioning; def., default protonation; ext. peptide, extended peptide; AF3, AlphaFold3. The value in the HMR column gives the integration timestep used after hydrogen mass repartitioning, or 2 fs where HMR was not applied. Coordinates were saved every 0.1 ns for all simulations. Force fields: a99SB-disp [52], a03w [211], DES-Amber [212].

**Table 4:**
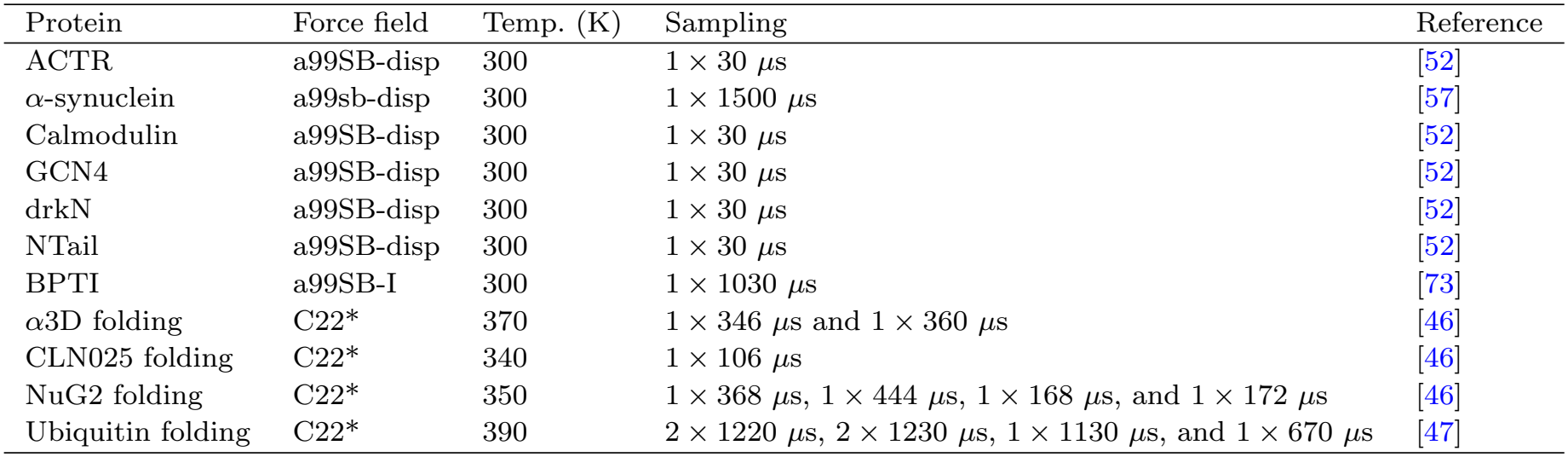
All-atom MD simulation datasets from the literature.

| Protein | Force field | Temp. (K) | Sampling | Reference |
|---|---|---|---|---|
| ACTR | a99SB-disp | 300 | 1 × 30 µs | [52] |
| α-synuclein | a99sb-disp | 300 | 1 × 1500 µs | [57] |
| Calmodulin | a99SB-disp | 300 | 1 × 30 µs | [52] |
| GCN4 | a99SB-disp | 300 | 1 × 30 µs | [52] |
| drkN | a99SB-disp | 300 | 1 × 30 µs | [52] |
| NTail | a99SB-disp | 300 | 1 × 30 µs | [52] |
| BPTI | a99SB-I | 300 | 1 × 1030 µs | [73] |
| α3D folding | C22* | 370 | 1 × 346 µs and 1 × 360 µs | [46] |
| CLN025 folding | C22* | 340 | 1 × 106 µs | [46] |
| NuG2 folding | C22* | 350 | 1 × 368 µs, 1 × 444 µs, 1 × 168 µs, and 1 × 172 µs | [46] |
| Ubiquitin folding | C22* | 390 | 2 × 1220 µs, 2 × 1230 µs, 1 × 1130 µs, and 1 × 670 µs | [47] |

### HyRes molecular dynamics simulations

Molecular dynamics simulations of *α*-synuclein, A*β*42, BecN, ChiZ, Δ131Δ, and Tau were performed using OpenMM v8.2 [89] with an optimised version of the original HyRes-GPU model, HyRes-Opt. A*β*42 and Tau were also simulated using the original HyRes-GPU model [60]. Simulations were carried out in the NVT ensemble in a cubic box of side length 200 nm with periodic boundary conditions, using a 4 fs time step and a Langevin thermostat with a friction coefficient of 0.01 ps^*−*1^. A total of 10,100 frames was saved at intervals of 120 ps for *α*-synuclein, 28 ps for A*β*42, 150 ps for BecN, 30 ps for ChiZ, 108 ps for Δ131Δ, and 1.17 ns for Tau, with the first 100 frames discarded as equilibration. The sampling intervals were chosen based on the empirical scaling relationship with sequence length, *N* : Δ*t*_save_ = max (28, 6 × 10^*−*3^*N* ^2^) ps, which ensures that the radius of gyration was sufficiently uncorrelated between consecutive saved frames. The temperature, ionic strength, and pH were set as detailed in Table 5. In HyRes-Opt, histidine is modeled with a fractional charge estimated using the Henderson–Hasselbalch equation assuming p*K*_*a*_ = 6, whereas it is neutral in HyRes-GPU. HyRes-Opt differs from HyRes-GPU in the Lennard-Jones *ϵ* parameters, the dielectric constant and cutoffs used to model salt-screened electrostatic interactions, and the parameters governing hydrogen bonds between backbone amide and carbonyl groups. All HyRes ensembles were converted to all-atom resolution models using cg2all [90] with the Mainchain model.

**Table 5:** HyRes MD simulation datasets of disordered proteins produced in this work. We sampled 10,000 uncorrelated frames for analysis (Methods). Force fields: HyRes-GPU [60], HyRes-Opt [35]

| Protein | Force field | Temp. (K) | Ionic strength (M) | pH |
|---|---|---|---|---|
| α-synuclein | HyRes-Opt | 288 | 0.058 | 7.4 |
| Aβ<sub>42</sub> | HyRes-GPU | 293 | 0.169 | 7.4 |
| Aβ<sub>42</sub> | HyRes-Opt | 293 | 0.169 | 7.4 |
| BecN | HyRes-Opt | 288 | 0.147 | 6.8 |
| ChiZ | HyRes-Opt | 298 | 0.069 | 7.0 |
| Δ131Δ | HyRes-Opt | 283 | 0.114 | 6.0 |
| Tau | HyRes-GPU | 293 | 0.169 | 7.4 |
| Tau | HyRes-Opt | 293 | 0.169 | 7.4 |

### Ensemble predictions with BioEmu and IDP-o

We used BioEmu to sample folding landscapes of small protein domains [4]. All predictions were made using version 1.1 of the model and default parameters. We reconstructed side chains using the built-in HPacker tool [91] followed by the default energy minimisation protocol. The proteins and number of conformations sampled with BioEmu are summarised in Table 6. For Tau we also used a 1000 frame ensemble predicted by the IDP-o model from previous work [11].

**Table 6:** Protein folding free energy landscapes sampled with BioEmu, number of conformations sampled and their folding free energies measured by orthogonal experiments (i.e., methods other than HDX). N.a., not applicable.

| **Protein** | **Samples (×1000)** | **Δ*G*<sub>fold</sub> (kcal mol<sup>−1</sup>)** |
|---|---|---|
| ACBP | 285 | -5.40 ± 0.50 [66] |
| ACP | 290 | -3.70 ± 0.40 [213] |
| *α*3D | 288 | -4.90 ± 0.50 [117] |
| *α*-synuclein | 4 | Disordered |
| CLN025 | 97 | -2.20 ± 0.10 [144] |
| GB3 | 291 | -3.92 ± 0.06 [214] |
| HP36 | 197 | -4.10 ± 0.30 [171] |
| Im7 | 587 | -6.30 ± 0.10 [63] |
| NuG2 | 295 | -7.63 ± 0.30 [29] |
| Ubiquitin | 977 | -6.83 ± 0.16 [34] |
| EEHEE-rd4-0642-6xHis | 71 | -3.25 ± 0.04 [62] |
| EEHEE-rd4-0642-No-His | 95 | -3.25 ± 0.04 [62] |
| EEHEE-rd4-0871-6xHis | 70 | -3.18 ± 0.04 [62] |
| EEHEE-rd4-0871-No-His | 96 | -3.18 ± 0.04 [62] |
| EEHEE-rd4-0871-K31L-E36V-No-His | 92 | n.a. |
| EHEE-rd2-0005-6xHis | 80 | -5.08 ± 0.05 [62] |
| HEEH-rd4-0097-6xHis | 81 | -3.30 ± 0.05 [62] |
| HHH-rd1-0012-6xHis | 82 | -3.17 ± 0.12 [62] |
| HHH-rd3-0062-6xHis | 85 | -3.61 ± 0.06 [62] |
| HHH-rd4-0518-6xHis | 74 | -4.61 ± 0.05 [62] |
| HHH-rd4-0518-No-His | 96 | -4.61 ± 0.05 [62] |
| HHH-rd4-0518-R35D-G45L-No-His | 98 | n.a. |
| HHH-rd4-0557-6xHis | 81 | -3.40 ± 0.05 [62] |
| LysM-0873-No-His | 98 | n.a. |
| LysM-1380-No-His | 99 | n.a. |
| LysM-3314-No-His | 99 | n.a. |

### Descriptors for protection factor forward models

Atomic distances, contacts, and per-atom solvent-accessible surface areas (SASA, using the Shrake–Rupley algorithm [92], a 0.14 nm probe radius and 960 sphere points) were calculated using MDTraj version 1.10.0 [93]. The front-facing atomic density, *ρ*_front_, was calculated for each backbone amide hydrogen by summing contributions from all heavy atoms within 0.60 nm. Each contribution was weighted by a smooth distance-switching function, [1 + (*r/*0.50 nm)^12^]^*−*1^, and an angular term, max(0, cos *θ*)^8^, where *θ* is the angle between the N–H bond vector and the vector from the amide hydrogen to the neighbouring atom. Lower angular exponents were also evaluated, as were calculations excluding heavy atoms belonging to residues *i* − 1 and *i* (Supplementary Figure 2d). These calculations were implemented in Python scripts utilising NumPy version 1.24.4 [94] and SciPy version 1.10.1 [95].

Electrostatic contributions were calculated using MDTraj version 1.10.0, OpenMM version 8.1.2 [89], and NumPy version 1.24.4. Atomic partial charges were assigned using the AMBER14SB force field [96]. For each backbone amide hydrogen *i*, the Debye–Hückel potential was calculated for every trajectory frame according to

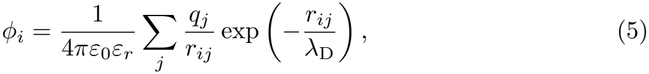

where *q*_*j*_ is the partial charge of atom *j, r*_*ij*_ is its distance from amide hydrogen *i*, and atoms belonging to residues *i*− 1 and *i* in the same chain were excluded. The Debye length was calculated as

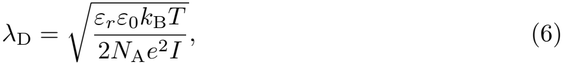

where *I* is the ionic strength in mol m^*−*3^. The temperature-dependent relative permittivity of water was given by the empirical relationship *ε*_*r*_ = 5321*/T* + 233.76 − 0.9297*T* + 1.417 × 10^*−*3^*T* ^2^ − 8.292 × 10^*−*7^*T* ^3^ [97]. Finally, the electrostatic contribution to the protection factor was calculated as

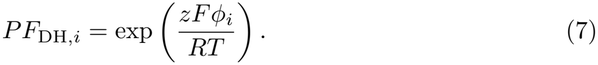

Here *z* = −1 is the charge of hydroxide, *F* is the Faraday constant, and *R* is the gas constant [41]. To approximately account for electronic polarisation effects, the Debye–Hückel calculation was additionally performed with all atomic partial charges uniformly scaled by a factor of 0.75. This model is referred to as DH_scaled.

### Explicit open-state models

Hydrogen exchange can be described using the Linderstrøm–Lang model, in which each backbone amide interconverts between closed, exchange-incompetent and open, exchange-competent conformations,

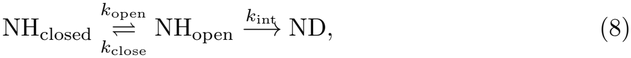

where *k*_open_ and *k*_close_ are the conformational opening and closing rates, respectively, and *k*_int_ is the intrinsic chemical exchange rate of the exposed amide. Applying the steady-state approximation to the open state gives

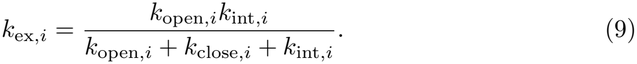

When conformational interconversion is fast compared with chemical exchange,

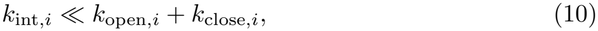

Eq. (9) reduces to

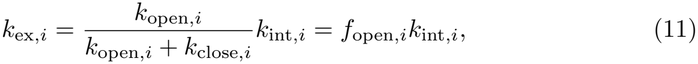

where the equilibrium open-state population corresponds to the structural ensemble as

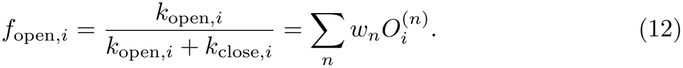

Here, 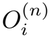 is the binary open-state assignment of amide *i* in conformation *n*, with 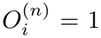 for an open conformation and 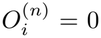 for a closed conformation, and *w*_*n*_ is the corresponding normalised ensemble weight of conformer *n*. The model-derived protection factor was therefore

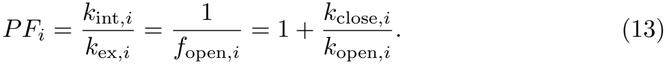

The reduction of Eq. (9) to Eq. (11) corresponds to the rapid-equilibration regime underlying EX2 exchange, in which conformational interconversion is fast relative to chemical exchange. More specifically, it requires *k*_int,*i*_ ≪ *k*_open,*i*_ +*k*_close,*i*_, which is more general than the commonly stated EX2 condition *k*_int,*i*_ ≪ *k*_close,*i*_. The frequently used approximation

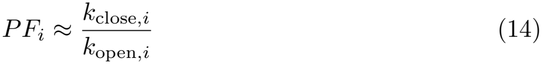

additionally requires *k*_open,*i*_ ≪ *k*_close,*i*_, such that the closed state is strongly populated. Equation (13) does not require this low-open-population assumption and therefore remains applicable to highly exposed amides in flexible or disordered regions. In the uncorrected binary open-state model, the limit *f*_open,*i*_ → 1 gives *PF*_*i*_→ 1, corresponding to exchange at the intrinsic rate.

Open and closed conformations were assigned using a binary classification. Closed conformations were assumed to be exchange-incompetent, whereas open conformations were assumed to exchange at the full intrinsic rate, except in model variants that explicitly accounted for modulation by the local electrostatic potential. Models M9– M14 differed only in the structural criteria used to define 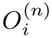.

In M9, a previously proposed model [23], an amide was classified as open when its polar-distance descriptor satisfied *d*_polar_ *>* 3.0 (minimum distance to a polar protein atom excluding residues *i* − 1 to *i*) and its solvent-accessible surface area (SASA) exceeded 4..0 Å^2^. The SASA threshold was selected by scanning possible values using the MD datasets of folded and disordered proteins (Supplementary Figure 2b, Tables 3 and 4). In M10, an amide was classified as open when it formed no hydrogen bond satisfying both a donor–acceptor distance below 0.26 nm and a hydrogen-bond angle above 120°, and when its amide-proton SASA exceeded 0.05 Å^2^. The low SASA threshold imposed a strict criterion for classifying an amide as buried and protected from exchange. M11 additionally required the directional packing descriptor *ρ*_front_ *<* 0.5, with atoms from residues *i* − 1 and *i* excluded from the calculation. This ensures that the path in front of the amide would be sterically accessible for a hydroxide to deprotonate the amide. M12 used the M10 criteria together with *ρ*_front_ *<* 0.5, without excluding atoms from neighbouring residues (i.e., amide protection can be induced by its own sidechain). M13 used the same hydrogen-bond criterion as M10 but applied a less stringent amide-proton SASA threshold of 0. 5 Å^2^ to define burial. M14 combined the M13 criteria with *ρ*_front_ *<* 0.5, again without excluding atoms from neighbouring residues. For all models, oxygen atoms belonging to residues *i* − 1 and *i* were excluded when evaluating hydrogen bonds involving amide *i*.

To account for residual protection observed in short peptides, the open-state populations calculated using M11, M12 and M14 were corrected according to

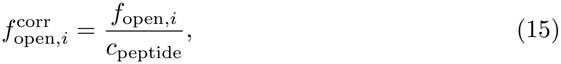

using *c*_peptide_ = 0.957319, 0.957089 and 0.951914 for M11, M12 and M14, respectively. These correction factors were obtained from MD simulations of Ace-AXA-NMe tripeptides (Table 3, Supplementary Figure 3b) by calculating the standard error-weighted mean PF of the central amides. The peptide correction factor, *c*_peptide_, was defined such that 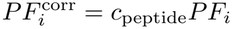, equivalently 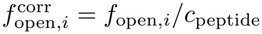, thereby normalising the mean predicted PF of the model peptides to unity.

We therefore define the conformation-specific effective exchange-competent observable

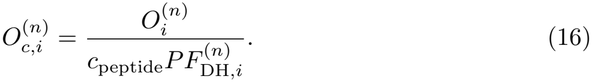

Here, 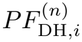 is the conformation-specific electrostatic contribution to the protection factor. For models without an electrostatic contribution, 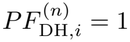, and for models without a peptide correction, *c*_peptide_ = 1.

The ensemble-averaged effective exchange-competent quantity is then

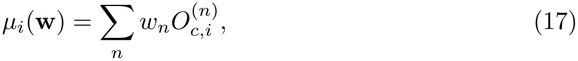

and the calculated protection factor is

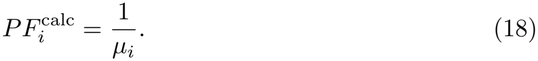

This formulation applies uniformly to the explicit open-state models with or without peptide and electrostatic corrections. Because the electrostatic contribution is included in 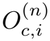 before ensemble averaging, correlations between structural opening and the local electrostatic environment are retained. This treatment assumes that solvent and ion distributions, and therefore the local hydroxide concentration, equilibrate more rapidly than the protein conformational dynamics.

These structural criteria and peptide correction factors for models M9–M14 are summarised in Table 7. We additionally compared these models with a previously proposed explicit-water model (Supplementary Figure 3), referred to here as the PH model [28]. In the PH model, an amide was classified as open when at least two water oxygen atoms were located within 0.26 nm of the amide hydrogen.

**Table 7:**
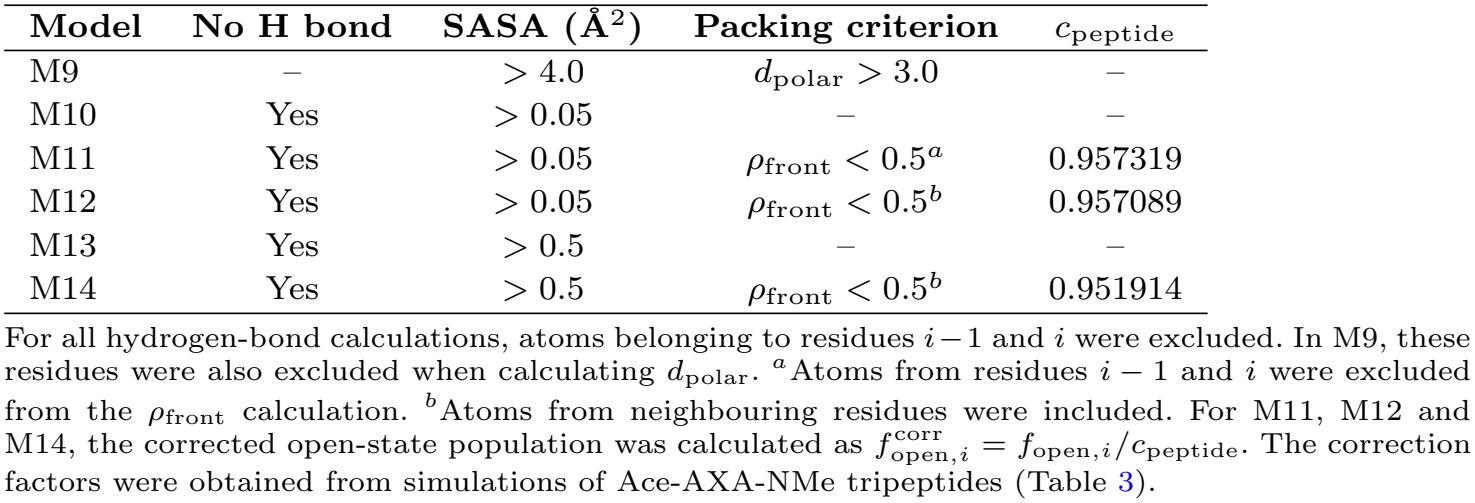
Structural criteria used to classify backbone amides as open (exchange-competent) in models M9–M14. SASA refers to the solvent-accessible surface area of the amide proton. A hydrogen bond was defined by a donor–acceptor distance below 0.26 nm and an angle above 120°.

| **Model** | **No H bond** | **SASA (Å<sup>2</sup>)** | **Packing criterion** | *c*<sub>peptide</sub> |
|---|---|---|---|---|
| M9 | – | > 4.0 | *d*<sub>polar</sub> > 3.0 | – |
| M10 | Yes | > 0.05 | – | – |
| M11 | Yes | > 0.05 | *ρ*<sub>front</sub> < 0.5<sup>*a*</sup> | 0.957319 |
| M12 | Yes | > 0.05 | *ρ*<sub>front</sub> < 0.5<sup>*b*</sup> | 0.957089 |
| M13 | Yes | > 0.5 | – | – |
| M14 | Yes | > 0.5 | *ρ*<sub>front</sub> < 0.5<sup>*b*</sup> | 0.951914 |

For all hydrogen-bond calculations, atoms belonging to residues *i* − 1 and *i* were excluded. In M9, these residues were also excluded when calculating *d*<sub>polar</sub>. <sup>*a*</sup>Atoms from residues *i* − 1 and *i* were excluded from the *ρ*<sub>front</sub> calculation. <sup>*b*</sup>Atoms from neighbouring residues were included. For M11, M12 and M14, the corrected open-state population was calculated as $f_{\mathrm{open},i}^{\mathrm{corr}} = f_{\mathrm{open},i}/c_{\mathrm{peptide}}$. The correction factors were obtained from simulations of Ace-AXA-NMe tripeptides (Table 3).

### Empirical forward models

We benchmarked our new forward models against previously proposed empirical forward models. The Best–Vendruscolo forward model (BV2006) [22] empirically estimates the protection factor of backbone amide *i* from local packing and hydrogen bonding:

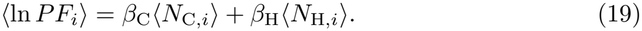

where *N*_C,*i*_ is the number of protein heavy atoms within a specified cutoff of the amide nitrogen, excluding atoms in residues *i* − 2 to *i* + 2, and *N*_H,*i*_ is the number of hydrogen-bond acceptors within 0.24 nm of the amide hydrogen. In the original BV2006 implementation, the contact cutoff was 0.65 nm and the parameters *β*_C_ = 0.35 and *β*_H_ = 2.0. We additionally implemented an angularly constrained variant, termed BV2006ang. In this model, contacts were counted within 0.50 nm and hydrogen bonds were required to satisfy both an H–acceptor distance cutoff of 0.24 nm and a donor– H–acceptor angle of at least 150°. The optimised coefficients were *β*_C_ = 0.89 and *β*_H_ = 4.39. These parameters were obtained by parameter scanning using the MD dataset (Tables 3, 4, 5) of folded and disordered states with experimentally determined NMR protection factors.

The M8 forward model [23] estimates the logarithmic protection factor of backbone amide *i* from its ensemble-averaged solvent-accessible surface area and distance to the nearest polar protein atom:

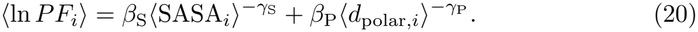

Here, SASA_*i*_ is the solvent-accessible surface area of the amide hydrogen and *d*_polar,*I*_ is its minimum distance to a protein N, O, S, or F atom, excluding atoms in residues *i*−1 to *i*+1. The fitted M8 parameters (using the MD dataset, Tables 3, 4, 5, of folded and disordered states) were *β*_S_ = 1.466, *γ*_S_ = 0.158, *β*_P_ = 46.840, and *γ*_P_ = 2.655.

### Ensemble reweighting with HX-NMR protection factors

Reweighting calculations with the BV2006 and BV2006ang models were performed with the BME software using ln(PF) values as restraints [32]. For the explicit open-state models, structural ensembles were reweighted against residue-specific HX-NMR protection factors using Bayesian maximum-entropy reweighting adapted to the non-linear relationship between the ensemble-averaged effective open-state quantity and the protection factor. Residues for which no exchange-competent conformation was sampled in the prior ensemble, such that 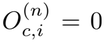 for all conformations and therefore *µ*_*i*_ = 0, were excluded from the reweighting because the corresponding calculated protection factor is infinite.

Using the notation introduced in Eq. (17), the calculated logarithmic protection factor was

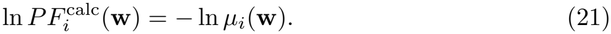

Protection factors were fitted in logarithmic space because they span several orders of magnitude and are naturally interpreted on a multiplicative scale. Fitting ln *PF* assigns residuals of equal magnitude to equivalent fold over- and underestimates, whereas fitting 1*/PF* directly in linear space disproportionately penalises ensembles that are too open. Experimental uncertainties were propagated according to

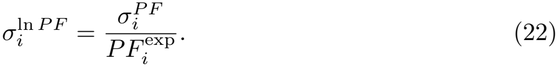

The posterior weights were obtained by minimising

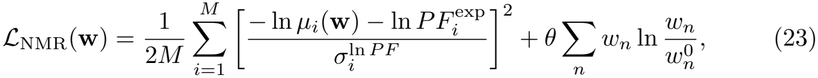

where *M* is the number of experimental protection factors, and 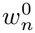 and *w*_*n*_ are the normalised prior and posterior conformational weights, respectively, with *w*_*n*_ ≥ 0 and ∑_*n*_ *w*_*n*_ = 1. The relative-entropy term corresponds to the Kullback–Leibler divergence,

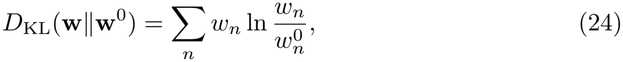

and *θ* controls the strength of this regularisation.

Defining

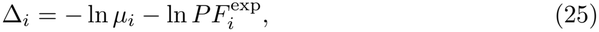

the gradient of the regularised objective with respect to conformational weight *w*_*n*_ is

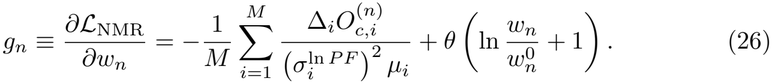

We implemented both direct optimisation of the conformational weights and a reduced-dimensional parametrisation, termed `log_lambda`. Direct weight-space optimisation used the prior-weighted softmax transformation

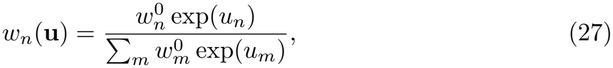

with

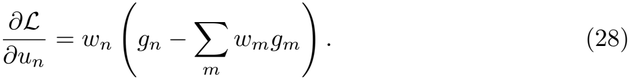

Motivated by Bayesian and maximum-entropy ensemble-refinement approaches, in which posterior weights can be expressed in an exponential Gibbs-family form and the optimisation recast in terms of a reduced set of generalised forces associated with the experimental observables [30–32], we additionally implemented a reduced-dimensional parametrisation of this form. In the present case, however, the fitted quantity ln 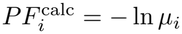 is a nonlinear function of the linearly ensemble-averaged quantity *µ*_*i*_. The resulting parametrisation is therefore motivated by, but is not identical to, the standard linear-observable Bayesian/maximum-entropy formulation.

In the `log_lambda` implementation, the posterior weights were parametrised as

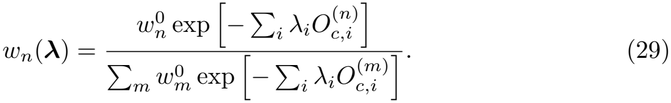

This reduces the number of optimisation variables from the number of conformations to the number of experimental observables and was therefore used by default. Because the fitted observable, ln 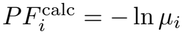, is nonlinear in the ensemble average, this is not the standard linear-observable BME dual formulation. Instead, the original loss in Eq. (23) is evaluated through the parametrised weights.

The derivation of the analytical gradients, the Gibbs-family form of Eq. (29), its relation to the stationary solution of the regularised objective, and numerical comparison with direct weight-space optimisation are described in Supplementary Note 1. Although global convexity is not formally guaranteed for the nonlinear ln *PF* objective, the two parametrisations yielded closely agreeing posterior ensembles across test systems and regularisation strengths tested. The loss and analytical gradients were minimised using L-BFGS-B over a range of *θ* values.

The effective fraction of the prior ensemble retained after reweighting was quantified from the relative entropy as

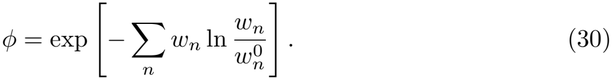

Reweighting was performed over a range of regularisation strengths *θ*, with each optimisation initialised from the prior ensemble. Posterior ensembles were selected based on the trade-off between agreement with experiment and deviation from the prior, as quantified by *ϕ*.

### Ensemble reweighting with HDX-MS data

For HDX-MS reweighting, the same ensemble-averaged effective open-state quantity,

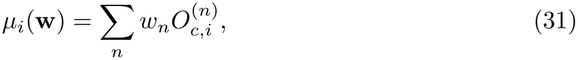

was used to predict residue-level exchange.

For peptide–time-point measurement *α*, measured after labelling time *t*_*α*_, the predicted fractional deuterium uptake was

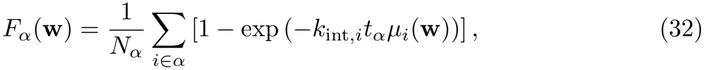

where *N*_*α*_ is the number of exchangeable backbone amides included for peptide *α*. The first two residues of each peptide and residues without an exchangeable backbone amide were excluded.

The posterior weights were obtained by minimising

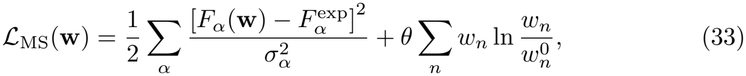

where *α* indexes all peptide–time-point measurements, and *F* ^exp^ and *σ*_*α*_ are the experimental mean fractional uptake and its uncertainty.

The gradient of the regularised objective with respect to conformational weight *w*_*n*_ is

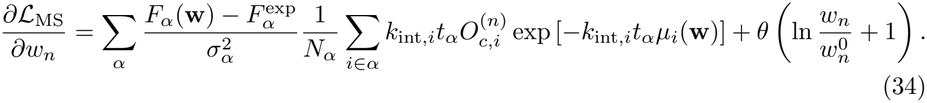

Because each HDX-MS measurement depends nonlinearly on multiple residue-level ensemble averages, and because overlapping peptides couple these residue-level quantities across measurements, HDX-MS reweighting was performed directly in the full weight space using a softmax parametrisation of the conformational weights. Each optimisation was initialised from the prior ensemble, and the relative-entropy regularisation was evaluated with respect to the corresponding prior weights. Gradients with respect to the unconstrained softmax variables were calculated using Eq. (28), with *g*_*n*_ = *∂ L*_MS_*/∂w*_*n*_. The detailed derivation of the gradient is provided in Supplementary Note 2. The loss and analytical gradient were minimised using L-BFGS-B over a range of regularisation strengths.

### Calculation of orthogonal data

Chemical shifts were predicted using SPARTA+ [98], and secondary chemical shifts were obtained using random-coil reference values calculated with POTENCI [99]. RDCs were calculated with PALES using a local alignment window of 15 residues [100]. PREs were predicted with DEER-PREdict using its MTSL spin-label rotamer library [101]. We assumed a transverse relaxation rate of 10 s^*−*1^ for the diamagnetic state and an effective spin-label correlation time, *τ*_*t*_, of 0.1 ns. SAXS profiles were calculated using Pepsi-SAXS [102], with the hydration-layer (*δρ*) and displaced-solvent (*r0*) parameters fixed to values determined previously [103].

Agreement between calculated and experimental chemical shifts was evaluated using a normalised RMSE, equivalent to the square root of the reduced chi-squared 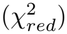 :

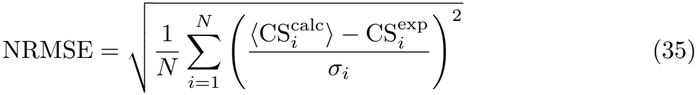

where *σ*_*i*_ is the nucleus-specific uncertainty of the calculated chemical shift arising from the forward model, as reported in the original SPARTA+ publication [98].

For RDCs, the magnitude of the ensemble-averaged calculated values was scaled to optimise agreement with experiment. The scaling factor was determined using residues for which the predicted RDCs had the same sign as the experimental values [55]. Agreement was quantified using the Q-factor calculated from the scaled RDCs:

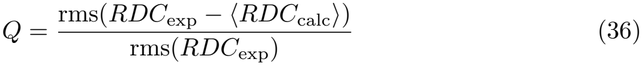

For PREs, agreement with experiment was quantified from the RMSE between the calculated and experimental intensity ratios 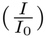. The correlation time of the nucleus-electron vector, *τ*_*C*_, was scanned from 1 to 20 ns in increments of 1 ns, and the value yielding the best agreement with the experimental data was selected.

The calculated SAXS scattering profiles were globally fitted to the experimental data using a multiplicative scale factor and a constant background. Least-squares fitting was performed using Scikit-learn [104], and agreement was quantified using the reduced 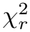 :

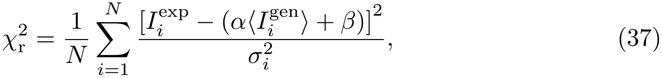

The experimental SAXS uncertainties were rescaled using the Bayesian Indirect Fourier Transform (BIFT) method [105].

All ensemble reweighting calculations using these types of orthogonal data were performed with the BME software [32]. Simultaneous reweighting with CS+SAXS and CS+PRE data was performed as previously described [55].

Amide N–H order parameters, *S*^2^, were calculated from rotationally aligned trajectories using the second moments of the N–H bond-vector orientations. Unit N–H vectors were divided into non-overlapping blocks with durations corresponding to 0.5, 1, 2, 3, 5, or 10 times the experimental isotropic rotational correlation time, *τ*_c_. For each block, the order parameter was calculated as

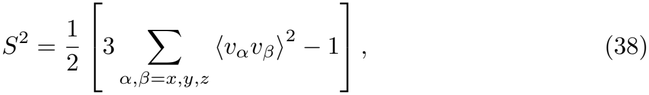

where *v*_*α*_ and *v*_*β*_ are the Cartesian components of the unit N–H bond vector and the angular brackets denote an average over frames within the block. To preserve the internal dynamics of the original trajectories, all frames within each block contributed uniformly to this average. HDX-derived frame weights were subsequently summed within each block, and the block-resolved *S*^2^ values were averaged using these summed weights to obtain the HX-reweighted ensemble values. For experimental datasets in which order parameters were derived using an effective N–H bond length of 1.02 Å, the calculated ensemble-averaged order parameters were subsequently scaled by a factor of 0.89 to account for zero-point vibrational averaging of the N–H bond vector.

Backbone three-bond scalar couplings, ^3^*J*, were calculated from the simulated backbone *ϕ* dihedral angles obtained using MDTraj, using Karplus relationships of the form ^3^*J* = *A* cos^2^(*ϕ* + *δ*) + *B* cos(*ϕ* + *δ*) + *C*. The parameters (*A, B, C, δ*) were (8.40, −1.36, 0.33, −60°) for ^3^*J*_HNHA_ [106], (3.71, −0.59, 0.08, 60°) for ^3^*J*_HNCB_ [106], (4.36, −1.08, −0.01, 180°) for ^3^*J*_HNCO_ [106], (4.41, −2.14, 0.77, 0°) for ^3^*J*_HACO_ [107], and (1.61, − 0.93, 0.55, 0°) for 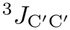 [108], with *A, B*, and *C* given in Hz. Couplings were calculated separately for every simulation frame, residue, and protein for which experimental data were available.

Nuclear Overhauser effect (NOE) distances were calculated for each trajectory frame using MDAnalysis version 2.4.3 [109]. For restraints involving equivalent or ambiguously assigned protons, all pairwise distances between the corresponding proton groups were included and combined using an *r*^*−*6^ average. Ensemble-averaged effective distances were calculated as

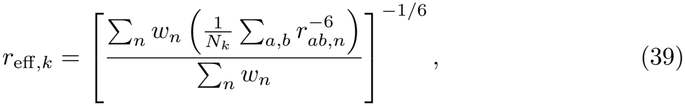

where *r*_*ab,n*_ is the distance between equivalent proton pair *a, b* for restraint *k* in frame *n, N*_*k*_ is the number of equivalent proton pairs, and *w*_*n*_ is the weight of frame *n*. Agreement with experiment was quantified from violations of the reported lower and upper distance bounds. Restraints with calculated distances within the experimental bounds were assigned zero error, whereas restraints outside the bounds contributed the magnitude of the corresponding distance violation. The NOE root-mean-square error was calculated over all restraints, including zero contributions from satisfied restraints.

To quantify changes in agreement with independent experimental data following HX-guided reweighting, we calculated the difference between the prior and posterior agreement metrics such that positive values indicated improved agreement and negative values indicated worsened agreement. Root-mean-square error (RMSE) was used for most orthogonal data types, while normalised RMSE was used for chemical shifts and the *Q*-factor for residual dipolar couplings (RDCs). For SAXS, we used the percentage change in reduced 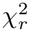 to account for differences in the absolute level of agreement between datasets. For each orthogonal data type, and separately for the IDP and folded-protein datasets, the resulting changes in predictive performance were normalised by the largest absolute change observed across all forward models and proteins within that data type and protein class (IDPs and folded proteins), yielding scores between 1 and 1. An overall cross-validation score for each forward model was then calculated by averaging these normalised scores across proteins and orthogonal data types.

### Analysis of folding free-energy landscapes

To characterise folding free-energy landscapes, we used the fraction of native contacts, *Q*, as a reaction coordinate. *Q* was calculated relative to the input PDB structure using a continuous switching function. Native contacts were defined as pairs of non-hydrogen atoms separated by at least four residues in sequence and by no more than 0.45 nm in the reference structure. Only contacts for which both residues were assigned as structured, i.e. non-coil, by DSSP [110] were included. Where present, an N-terminal hexahistidine tag and the three following residues were excluded from the calculation. For each trajectory frame, *Q* was calculated as

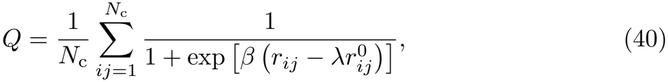

where *N*_c_ is the total number of native contacts, *r*_*ij*_ is the instantaneous distance between atoms *i* and *j*, 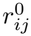 is the corresponding distance in the reference structure, *r*^0^ *λ* = 1.8, and *β* = 50 nm^*−*1^.

Folding free energies were calculated from the relative populations of the folded and unfolded states along *Q*. The boundary between these states was defined as the position of the maximum in the prior (i.e., before ensemble reweighting with experimental data) free-energy profile between *Q* = 0.2 and *Q* = 0.8, corresponding to the principal free-energy barrier separating the native basin from lower-*Q* conformations. Frames with *Q* above this cutoff were assigned to the folded state, whereas frames with *Q* below the cutoff were assigned to the unfolded state. The folding free energy was then calculated as

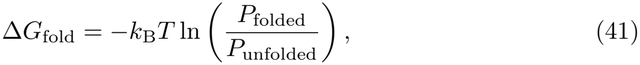

where *P*_folded_ and *P*_unfolded_ are the populations of the corresponding states. All folding free energies were evaluated at *T* = 298 K and reported in kcal mol^*−*1^. Backbone root-mean-square deviations (RMSDs) were calculated for folded proteins using GROMACS 2024. Trajectory frames were first fitted to the corresponding reference structure by least-squares superposition of the backbone atoms, and the RMSD was then calculated over the same atom selection.

The radius of gyration, *R*_g_, was calculated for each trajectory frame using MDTraj. All protein atoms, including terminal capping groups, were included, and atomic masses were used in the calculation.

## Data availability

Molecular dynamics ensembles of the intrinsically disordered proteins are available on Zenodo at https://zenodo.org/records/23569858. Molecular dynamics data of all folded proteins are available on Zenodo at https://zenodo.org/records/23569607 and https://zenodo.org/records/23568205. Peptide simulation trajectories including water coordinates are available on Zenodo at https://zenodo.org/records/23539119. The previously reported molecular dynamics trajectories (Table 4) are available for non-commercial use upon request from D. E. Shaw Research. All BioEmu-generated ensembles are available on Zenodo at https://zenodo.org/records/23564156. All previously published experimental data used for ensemble reweighting and validation are available on GitHub (https://github.com/KULL-Centre/2026_streit_hx). Millisecond HDX-MS data of the disordered proteins presented in this study will be available on PRIDE (https://www.ebi.ac.uk/pride) with the accession code PXD086174.

## Code availability

The new forward models and reweighting methods have been implemented in an open-source Python package, HXEnsemble, available on GitHub (https://github.com/KULL-Centre/HXEnsemble). All custom code, scripts and Jupyter notebooks used for the analyses presented in this work are also available on GitHub (https://github.com/KULL-Centre/_2026_streit_hx).

## Acknowledgments

We thank Susan Marqusee for discussions on interpretations of hydrogen exchange experiments. This work was sponsored by Peptone Ltd. J.O.S. acknowledges support by the European Molecular Biology Organisation through Postdoctoral Fellowship grant ALTF 98-2025. We acknowledge access to computational resources from the ROBUST Resource for Biomolecular Simulations (supported by the Novo Nordisk Foundation grant no. NNF18OC0032608) and via a grant from the Carlsberg Foudation (CF21-0392). We also acknowledge use of the Baskerville Tier 2 HPC service (https://www.baskerville.ac.uk/), which was funded by the EPSRC and UKRI through the World Class Labs scheme (EP/T022221/1) and the Digital Research Infrastructure programme (EP/W032244/1) and is operated by Advanced Research Computing at the University of Birmingham. The research was also supported by the PRISM (Protein Interactions and Stability in Medicine and Genomics) centre funded by the Novo Nordisk Foundation (NNF18OC0033950, to K.L.-L.). We thank D. E. Shaw Research for sharing previously published MD trajectories.

## Author contributions

J.O.S., M.I., S.B., and K.L-L. conceived and designed the study. J.O.S., K.T. and K.L-L. sourced the funding. J.O.S. performed all-atom MD simulations, collected the literature data, developed the forward models, implemented the software, reweighted, and analysed all conformational ensembles with input from M.I., S.B., and K.L-L. D.P. performed and analysed the millisecond HDX-MS experiments with input from P.K. G.T. performed simulations with the HyRes force fields. T.K.S. contributed to the collection of the literature data. J.O.S wrote the initial draft of the manuscript and all authors contributed to the editing and reviewing of the manuscript.

## Supplementary Information

### Supplementary Notes

#### Supplementary Note 1: Reweighting with HX-NMR protection factors

For the explicit open-state models, each conformation contributes an effective exchange-competent observable 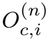 for residue *i*, defined in Eq. (16). Under ensemble weights **w**,

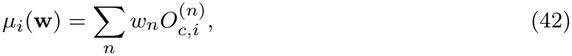

and the calculated logarithmic protection factor is

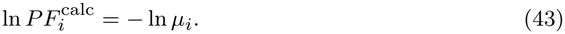

Thus, the effective exchange-competent observable is averaged linearly over the ensemble before conversion to a protection factor or logarithmic protection factor.

For the data term,

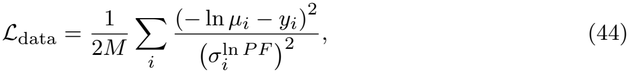

where 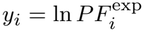 and 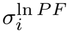 is the corresponding uncertainty in logarithmic protection-factor space. Defining

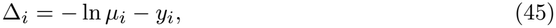

and noting that

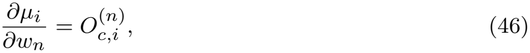

the derivative of the data term is

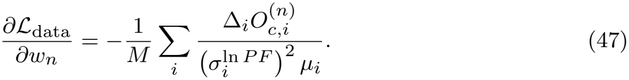

Including the relative-entropy term gives

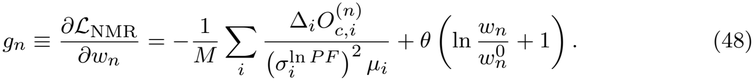

For direct optimisation in unconstrained softmax variables,

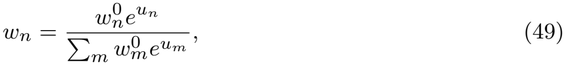

the corresponding gradient is

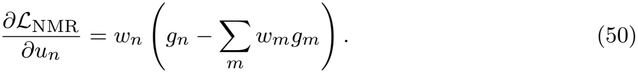

The origin of the Gibbs-family parametrisation can be seen from the first-order optimality condition of the original regularised objective with respect to the conformational weights, subject to the normalisation constraint ∑_*n*_ *w*_*n*_ = 1. Introducing a Lagrange multiplier *ξ* for this constraint gives

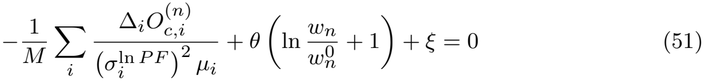

at a stationary point. Rearranging,

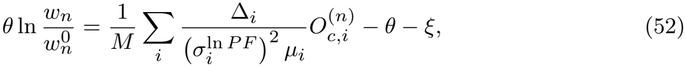

and therefore

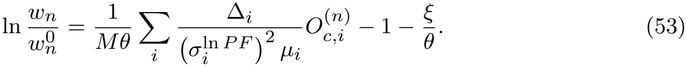

Exponentiating gives

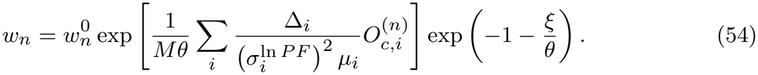

The final factor is independent of *n* and therefore contributes only to the normalisation. Thus,

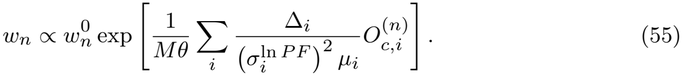

Defining

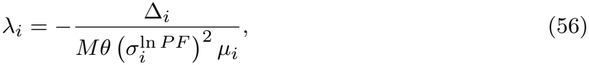

the stationary weights can be written in the Gibbs-family form

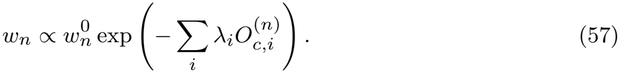

This motivates the reduced `log_lambda` parametrisation used in the main text. Importantly, this relation applies at a stationary solution because both Δ_*i*_ and *µ*_*i*_ depend on the posterior weights. The reduced parametrisation is therefore motivated by the Gibbs-family structure necessarily satisfied by stationary solutions of the original regularised weight-space objective.

Differentiating the normalised Gibbs weights gives

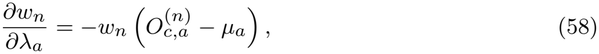

where

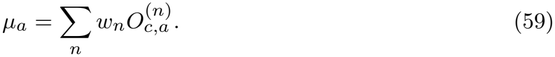

The gradient with respect to the multiplier *λ*_*a*_ is therefore

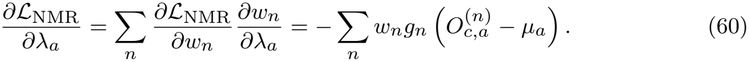

The `log_lambda` parametrisation reduces the dimensionality of the optimisation from the number of conformations to the number of experimental observables. It differs, however, from the conventional BME dual formulation for linear observables because the fitted quantity *−* ln *µ*_*i*_ depends nonlinearly on the ensemble average. The multipliers are therefore related self-consistently to the residuals at the optimum rather than being obtained from the standard closed-form linear-observable BME dual problem.

Accordingly, global convexity of the objective is not formally guaranteed. For the contribution of a single protection factor to the data term,

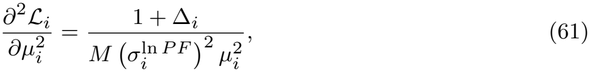

which can become negative when Δ_*i*_ *< −*1. The relative-entropy term is convex in the ensemble weights and regularises the optimisation, but does not provide a general guarantee of global convexity for arbitrary values of *θ*.

For each value of *θ*, optimisation was initialised from the defined prior ensemble. The effective fraction of the prior ensemble retained after reweighting was quantified from the relative entropy as

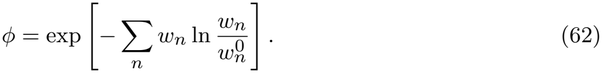

Reweighting was performed over a range of *θ* values to generate a trade-off between agreement with the HX data and deviation from the prior ensemble.

As a numerical validation of the reduced parametrisation, `log_lambda` optimisation was compared with direct optimisation in the full softmax weight space, with both calculations initialised from the same prior ensemble. The two implementations yielded closely agreeing posterior ensembles across the systems and regularisation strengths tested, with Kullback– Leibler divergences between the resulting posterior weight distributions typically below 10^*−*4^ and not exceeding 0.03 across all cases tested. The largest discrepancies were observed under strong reweighting, corresponding to small *θ* and low effective ensemble fractions, where the high-dimensional weight-space optimisation was comparatively more sensitive to convergence tolerance.

Protection factors are ensemble-averaged observables and therefore do not uniquely determine all microscopic degrees of freedom of a conformational ensemble. The purpose of the regularised reweighting is consequently not to infer a unique microscopic ensemble, but to identify a posterior ensemble that improves agreement with the HX data while remaining minimally perturbed from the defined prior. The physical relevance of the resulting posterior ensembles was assessed throughout this work by cross-validation against orthogonal experimental observables.

#### Supplementary Note 2: Reweighting with HDX-MS uptake data

For peptide *α* at labelling time *t*_*α*_, the predicted fractional uptake is

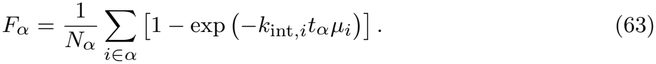

Defining

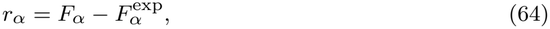

the HDX-MS data term is

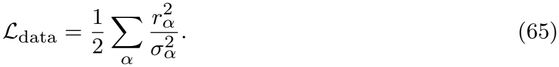

Since

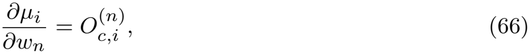

the derivative of the peptide uptake is

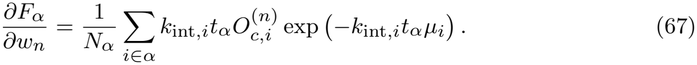

The complete gradient is therefore

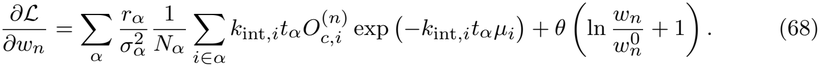

The corresponding gradient in unconstrained softmax variables follows from Eq. (28). Because each HDX-MS measurement depends nonlinearly on several residue-level ensemble averages, reweighting was performed directly in the full softmax weight-space parametrisation. As for HX-NMR reweighting, optimisation was performed over a range of *θ* values and initialised from the prior ensemble, with *ϕ* used to quantify the extent of reweighting.

## Supplementary Figures

**Supplementary Figure 1.**
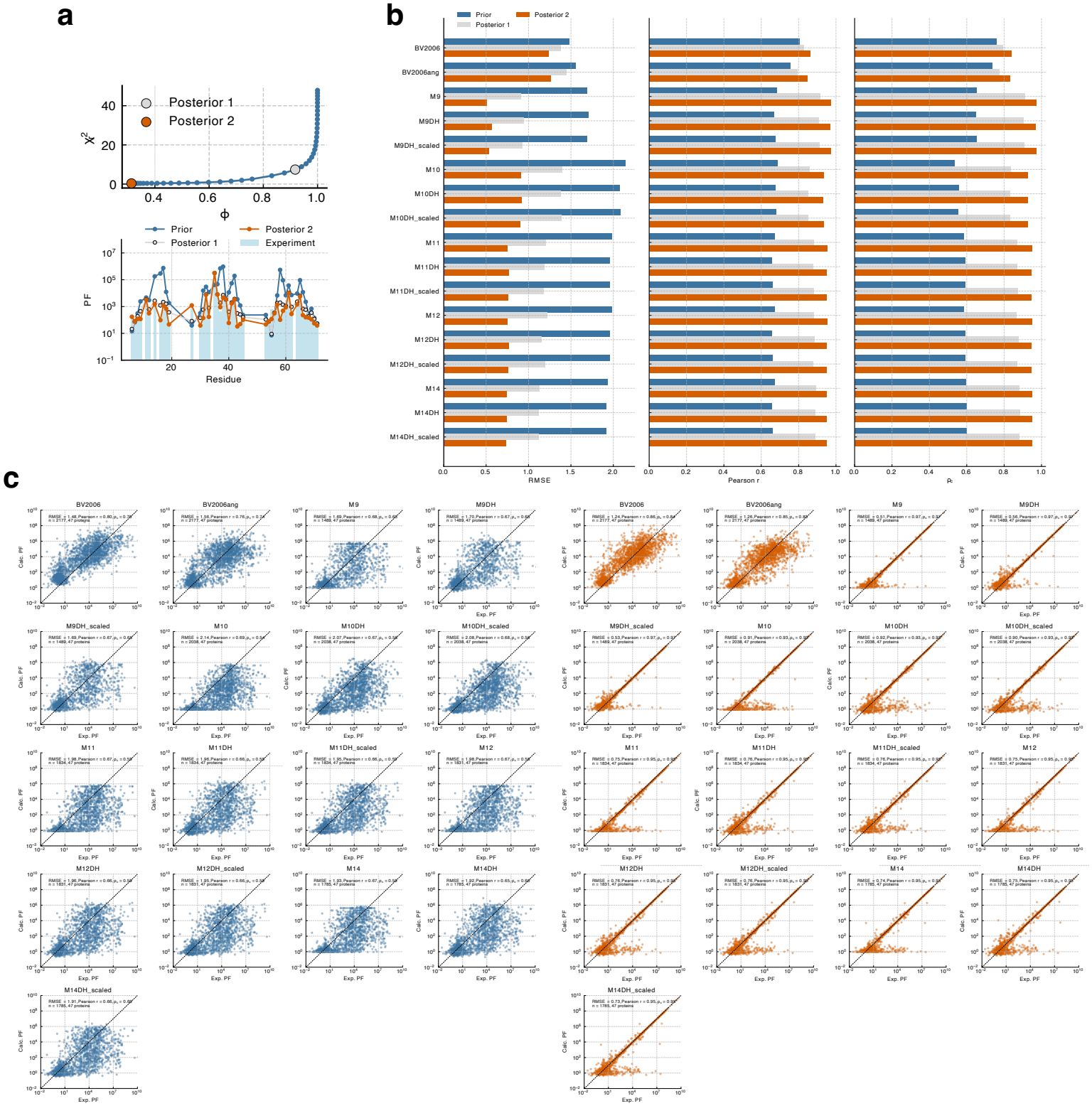
Assessment of forward models with MD ensembles before and after reweighting with HDX-NMR data. **a**. Example reweighting L-curve for an MD ensemble of *α*3*D*, showing the reduced 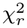 as a function of the effective fraction of frames, *ϕ*. The elbow defines the posterior 1 ensemble, while the final reweighted solution (posterior 2) is selected as the solution with the lowest 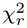 subject to *ϕ >* 0.1. Below, HDX-NMR protection factors are compared with predictions from the prior and reweighted ensembles. **b**. Performance of the tested forward models quantified in log_10_ space by the root-mean-square error (RMSE), Pearson correlation coefficient (*r*), and concordance correlation coefficient (*ρ*_*c*_) using the prior ensembles and after reweighting. posterior 1 corresponds to the elbow of the optimization L-curve, whereas posterior 2 corresponds to a minimum fraction of contributing frames of *ϕ* = 0.1. **c**. Correlation between experimental (Exp.) and calculated (Calc.) PFs across amides and proteins in the dataset. Predictions are shown for the prior ensembles (left) and after ensemble reweighting (posterior 2; right). All 2,177 amides were included for the empirical models, whereas analyses using explicit models were restricted to amides that sampled an open state at least once under the respective model definition.

**Supplementary Figure 2.**
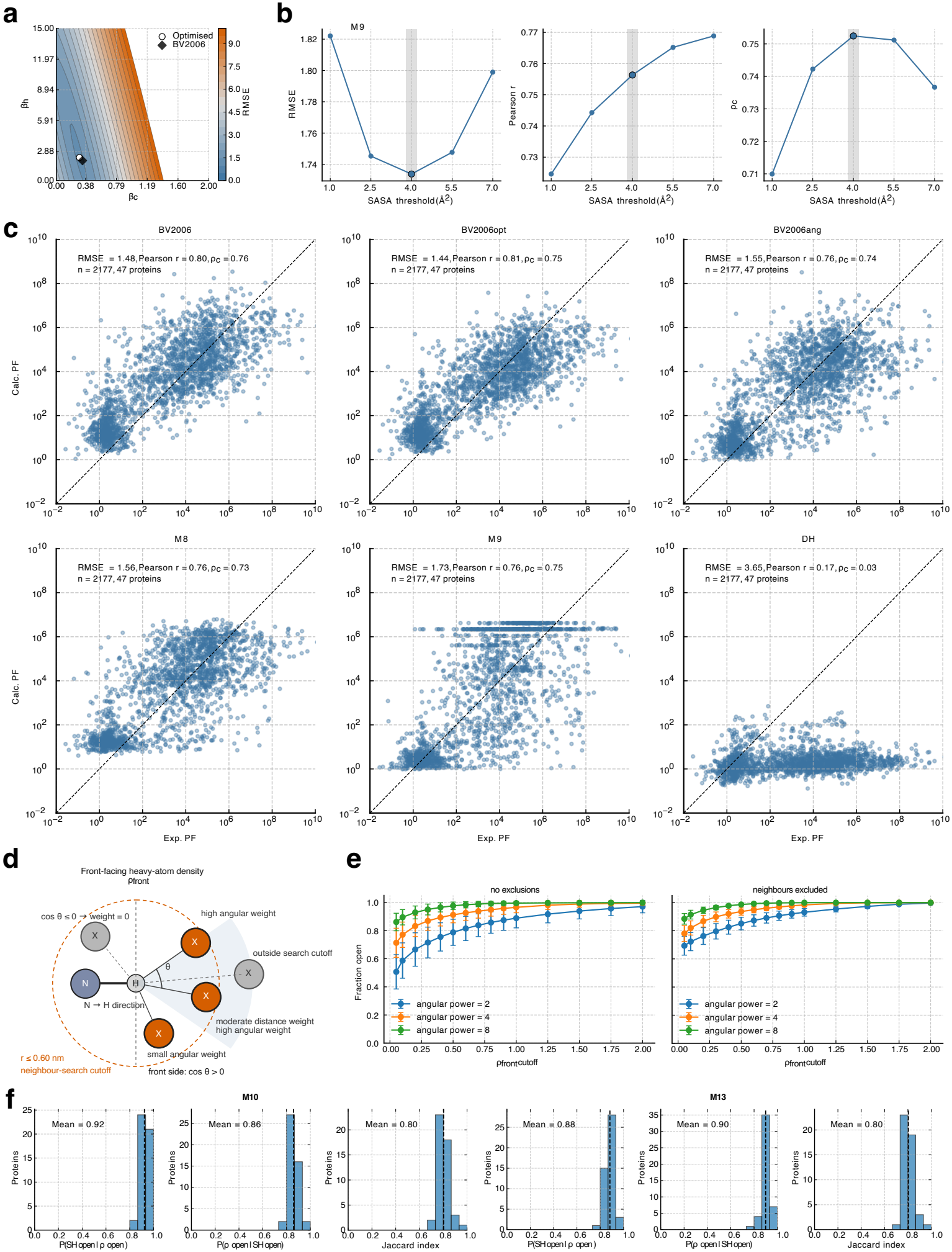
Analyses of additional forward models and structural descriptors for hydrogen exchange. **a**. Optimisation of BV2006 empirical parameters on the MD dataset using RMSE, yielding values close to those reported previously. **b**. Scan of the M9 solvent-accessible surface area (SASA) cutoff on the MD dataset using RMSE and *ρ*_*c*_, identifying 4.0 Å^2^ as the optimal cutoff. **c**. Correlation between experimental (Exp.) and calculated (Calc.) PFs across the MD dataset for additional forward models without reweighting. BV2006opt uses the optimised parameters from panel a, BV2006ang includes an angular hydrogen-bond criterion, M8 uses an alternative empirical functional form, and DH includes only the Debye-Hückel electrostatic contribution (Methods). **d**. Schematic of the *ρ*_front_ descriptor quantifying atomic packing in front of the amide N–H bond vector. The angle *θ* determines the directional weighting of neighbouring atoms applied to a contact switching function (Methods). **e**. Fraction of open amide conformations (mean *±* s.d.) from MD simulations of 19 non-proline Ace–AXA–NMe tripeptides across *ρ*_front_ cutoffs and angular powers (cos(*θ*)^*power*^), without (left) and with (right) exclusion of neighbouring residues. **f**. Redundancy between open conformations identified by SASA and hydrogen-bond criteria (models M10 and M13, with SASA cutoffs of 0.05 Å^2^ and 0.5 Å^2^, respectively) and *ρ*_front_ across the MD dataset, quantified by the Jaccard index.

**Supplementary Figure 3.**
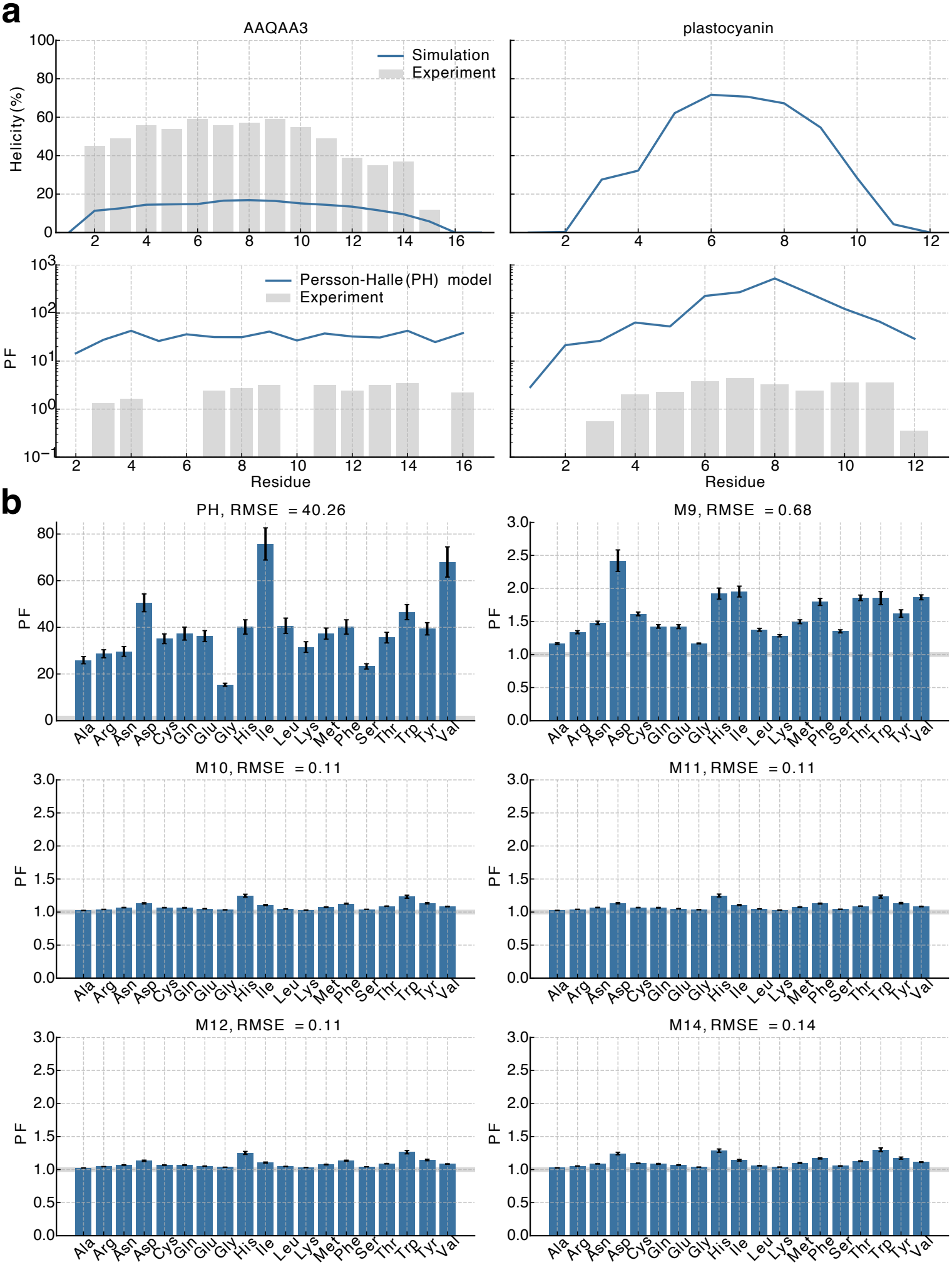
Comparison of explicit open-state models on disordered peptides and tripeptides. **a**. Helical propensities (top) from MD simulations and experiment for (AAQAA)_3_, and PFs (bottom) predicted with the Persson–Halle (PH) explicit-water model and compared with experiment. **b**. Predicted PFs for all 19 non-proline Ace–AXA–NMe tripeptides (mean *±* s.e. from blocking) using different explicit open-state models. RMSE values quantify deviations from the expected base-line of PF = 1.

**Supplementary Figure 4.**
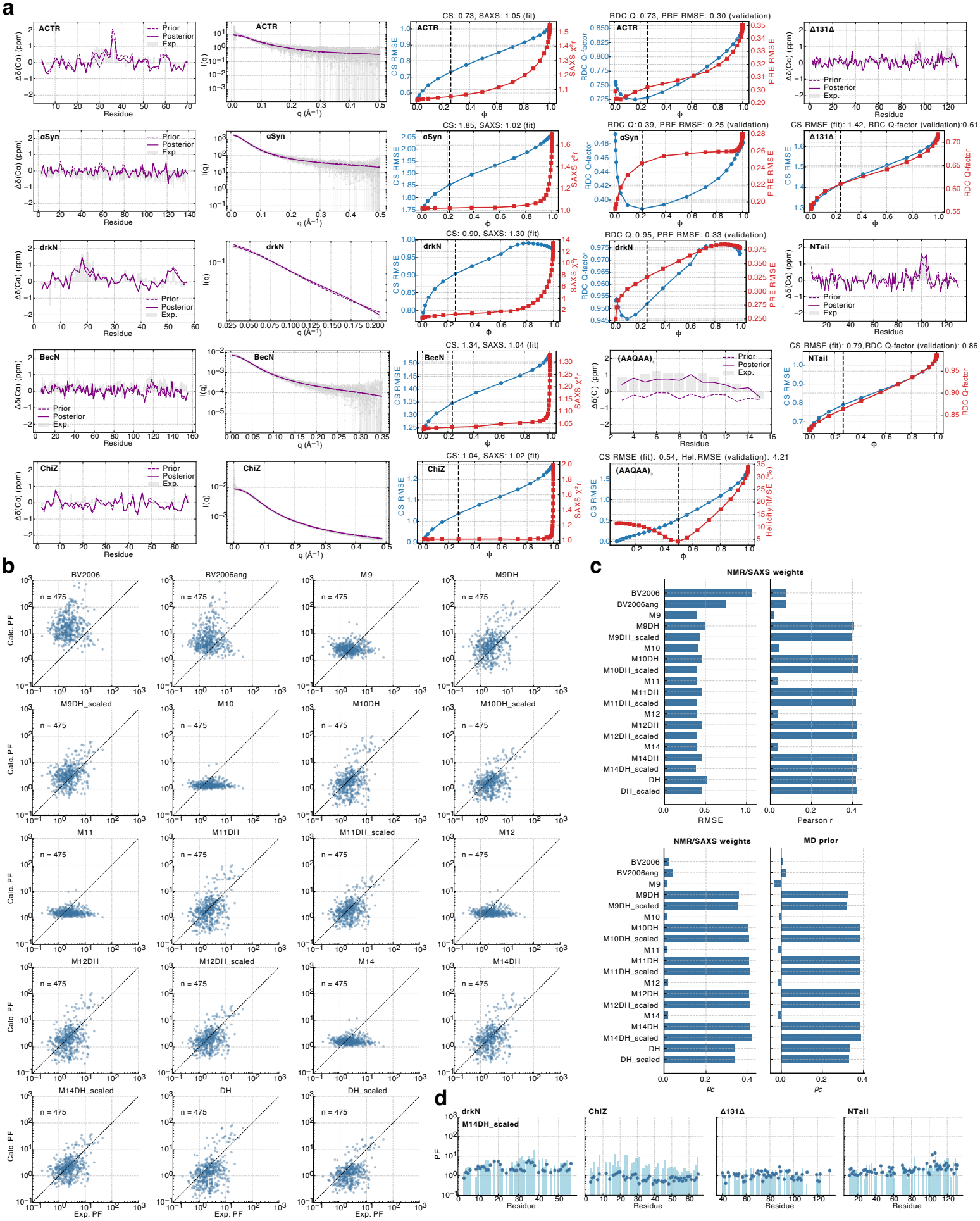
Reweighting of disordered protein ensembles with NMR and SAXS data and benchmarking of HX forward models on the refined ensembles. All ensembles were generated using all-atom MD simulations, except those of the IDPs *α*-synuclein, BecN, ChiZ, Δ131Δ, which were simulated using the HyRes-Opt force field (Tables 3– 5). **a**. Agreement with NMR chemical shifts (representative C*α* secondary shifts shown) and SAXS profiles before (prior) and after (posterior) reweighting for ACTR, *α*-synuclein, BecN, ChiZ, and drkN (left), with reweighting curves showing chemical shift and SAXS agreement as a function of effective ensemble size (*ϕ*; middle). ACTR, *α*-synuclein, and drkN were cross-validated against RDC and PRE data (right). (AAQAA)_3_, Δ131Δ, and NTail were reweighted using backbone chemical shifts and cross-validated against helical propensities derived from chemical-shift temperature dependencies ((AAQAA)_3_) or RDCs (Δ131Δ and NTail). **b**. Correlation between experimental (Exp.) and calculated (Calc.) PFs for 475 amides from experimentally refined disordered protein ensembles. **c**. RMSE, Pearson correlation coefficient, and *ρ*_*c*_ of forward models using the experimentally refined ensembles, with *ρ*_*c*_ for the corresponding MD priors with uniform weights shown for comparison. **d**. Experimental and M14DH_scaled-predicted PFs for four representative disordered proteins not shown in the main text.

**Supplementary Figure 5.**
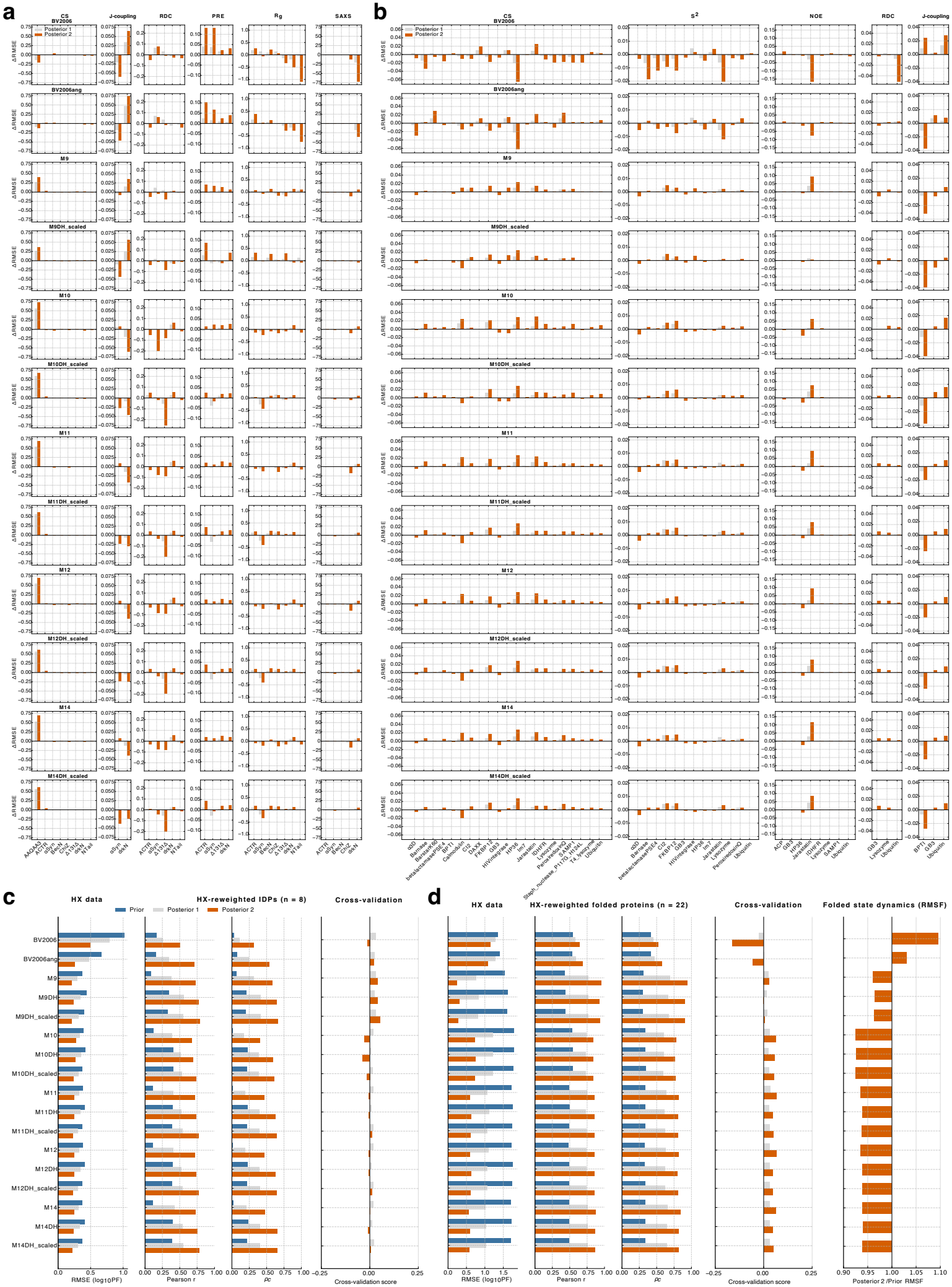
Cross-validation of HX-reweighted MD ensembles of disordered and folded proteins. All ensembles were generated using all-atom MD simulations, except those of the IDPs *α*-synuclein, BecN, ChiZ, Δ131Δ, which were simulated using the HyRes-Opt force field (Tables 3– 5). **a–b**. Change in agreement between prior and posterior ensembles for disordered (a) and folded (b) proteins after reweighting with HX-NMR data, shown for different orthogonal data types (columns) and forward models (rows). Positive values indicate improved and negative values worsened agreement. Changes are reported as prior–posterior normalised RMSE for chemical shifts; RMSE for *J*-couplings, PRE intensity ratios, *R*_*g*_, *S*^2^ order parameters, and NOE distance-restraint violations; Q-factor for RDCs; and percentage change in reduced 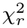 for SAXS (Methods). **c**. Forward-model performance for HX data (RMSE, Pearson correlation coefficient, and *ρ*_*c*_; left) and cross-validation against orthogonal data (right) for disordered proteins. Cross-validation scores are averaged across proteins and orthogonal data types, with positive and negative values indicating improved and worsened agreement, respectively. **d**. Equivalent analysis for folded proteins, together with the average change in backbone flexibility after HX reweighting (posterior 2/Prior RMSF). Values below and above 1 indicate decreased and increased flexibility, respectively.

**Supplementary Figure 6.**
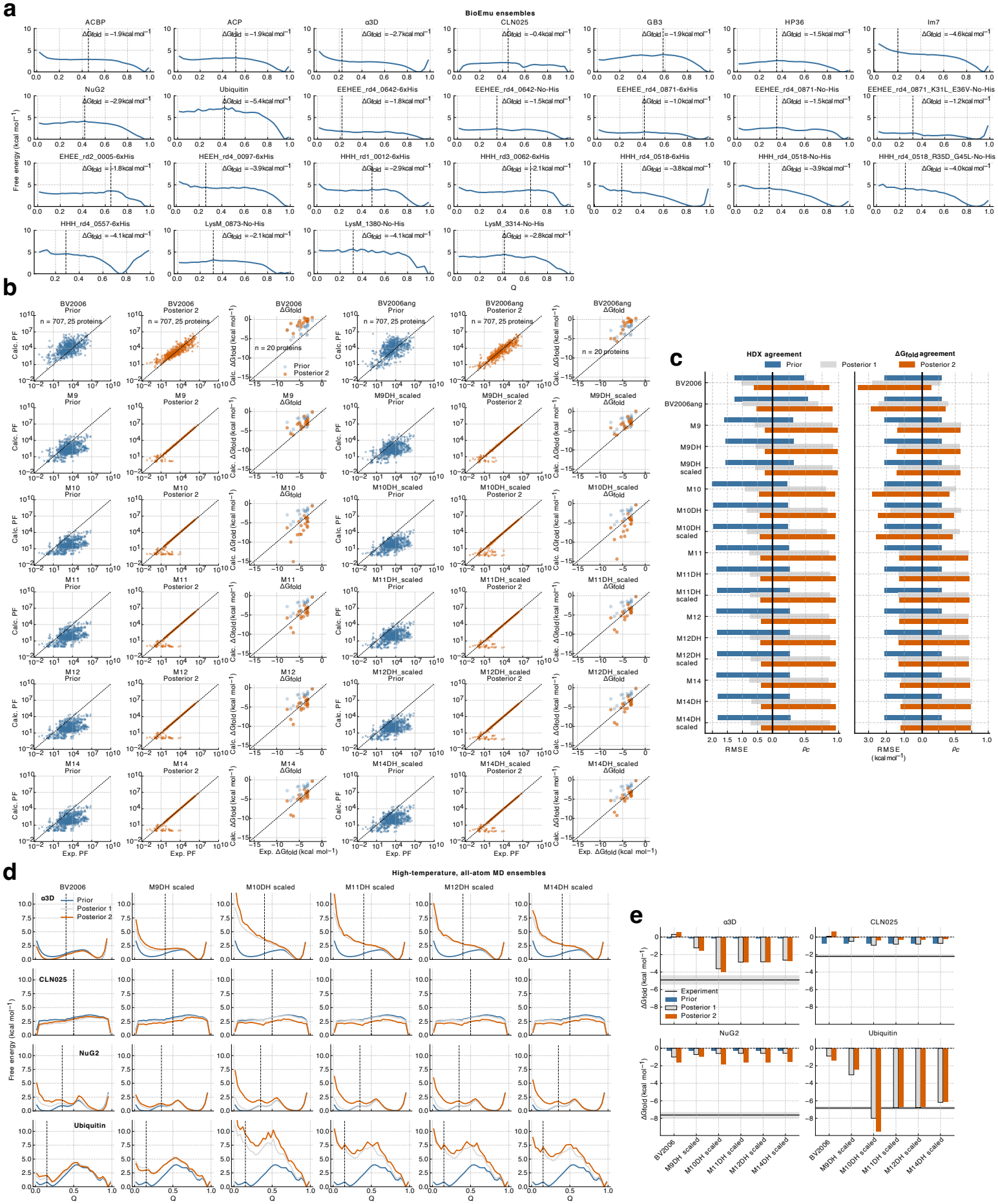
HDX-NMR reweighting and cross-validation of protein folding free-energy landscapes. **a**. Folding free-energy landscapes as a function of the fraction of native contacts (*Q*) for all 25 proteins generated with BioEmu before HDX reweighting. The vertical line indicates the cutoff used to separate unfolded and folded conformations, and the corresponding calculated folding free energy (Δ*G*_fold_) is shown. **b**. Correlation between experimental (Exp.) and calculated (Calc.) PFs across 707 amides from 25 proteins for different forward models, shown for the prior and posterior 2 ensembles. Agreement between experimental and calculated Δ*G*_fold_ values is shown for the subset of 20 proteins with orthogonally measured folding stabilities. **c**. Forward-model performance for HDX data and cross-validation against experimental Δ*G*_fold_ values from panel b before and after reweighting. **d**. Folding free-energy landscapes of four proteins obtained from previously published all-atom, millisecond-timescale reversible folding simulations near their melting temperatures, shown before and after HDX-NMR reweighting with different forward models. The vertical line indicates the cutoff used to separate unfolded and folded conformations for calculation of Δ*G*_fold_. For this ubiquitin ensemble, a cutoff of *Q* = 0.15 was used to exclude a partially folded state spanning approximately *Q* = 0.15–0.4 that retains substantial *β*-strand structure. **e**. Calculated (bars) and experimental (horizontal line) Δ*G*_fold_ values for the four proteins in panel d, shown for different forward models before and after reweighting.

**Supplementary Figure 7.**
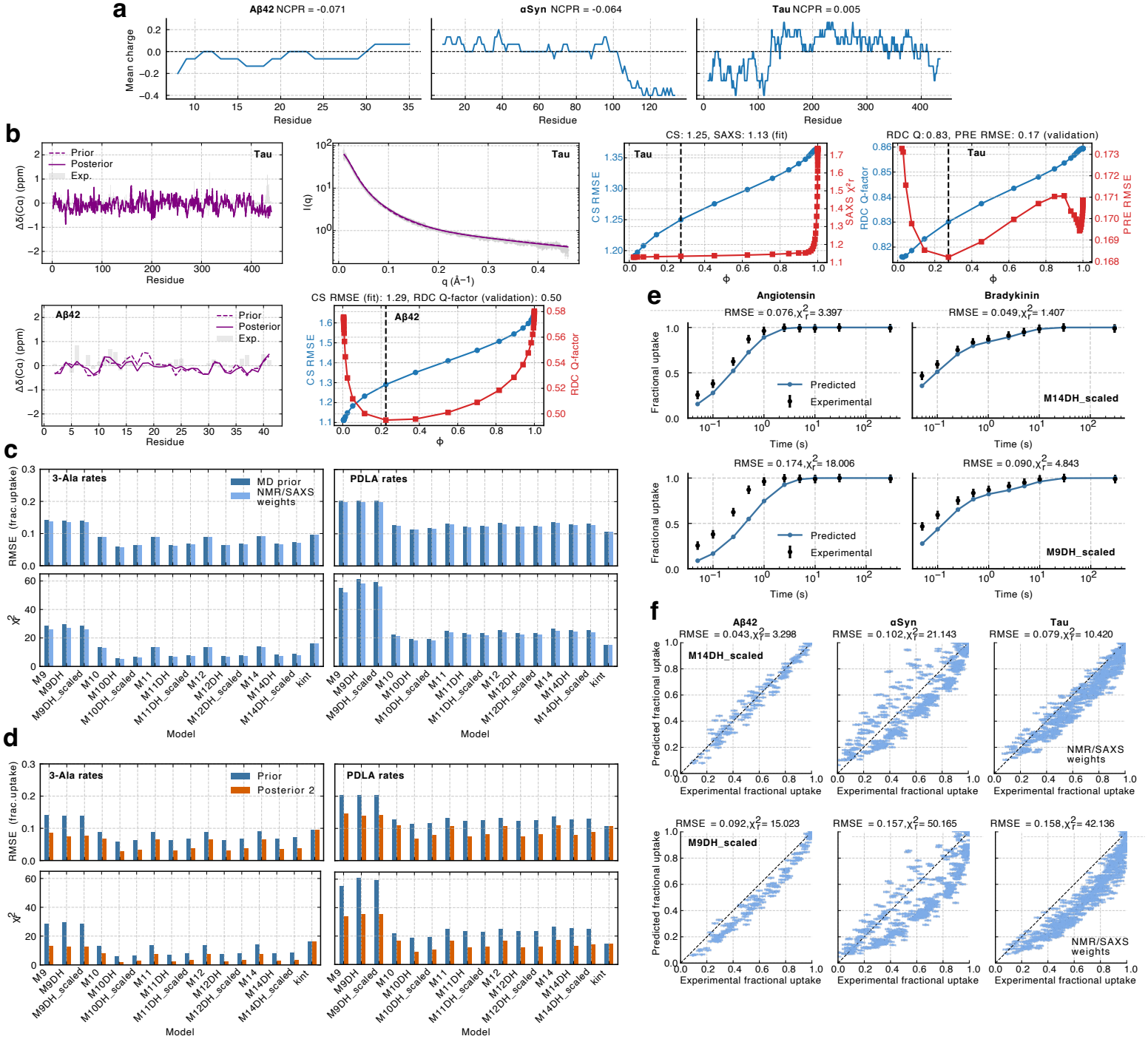
Millisecond HDX-MS of disordered proteins and peptides for bench-marking HDX-MS forward models. **a**. Net charge per residue (NCPR) and mean charge along the sequence, averaged over a 15-residue sliding window, for the three intrinsically disordered proteins (IDPs) studied here. **b**. Reweighting (fit) and cross-validation (validation) of Tau (top) and A*β*42 (bottom) ensembles. NMR chemical shifts and SAXS data were used to reweight Tau, with RDC and PRE data used for cross-validation. The A*β*42 ensemble was reweighted using NMR chemical shifts and cross-validated against RDCs. **c**. Performance of forward models in predicting HDX-MS fractional uptake across the three IDPs and two peptides (angiotensin and bradykinin), quantified by RMSE and reduced 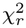. Results are shown using 3-Ala and PDLA intrinsic reference rates, and for MD prior ensembles with uniform weights and experimentally refined ensembles of the IDPs (NMR/SAXS weights). **d**. Equivalent benchmark to panel c before and after HDX-MS reweighting, using posterior 2 with a minimum effective fraction of frames of *ϕ >* 0.1. **e**. Experimental and predicted fast HDX-MS uptake curves for angiotensin and bradykinin using the uniform MD prior, 3-Ala reference rates, and two representative forward models. **f**. Correlation between experimental and predicted HDX-MS fractional uptake values for all peptides from the three IDPs using experimentally refined ensembles (NMR/SAXS weights), 3-Ala reference rates, and two representative forward models.

**Supplementary Figure 8.**
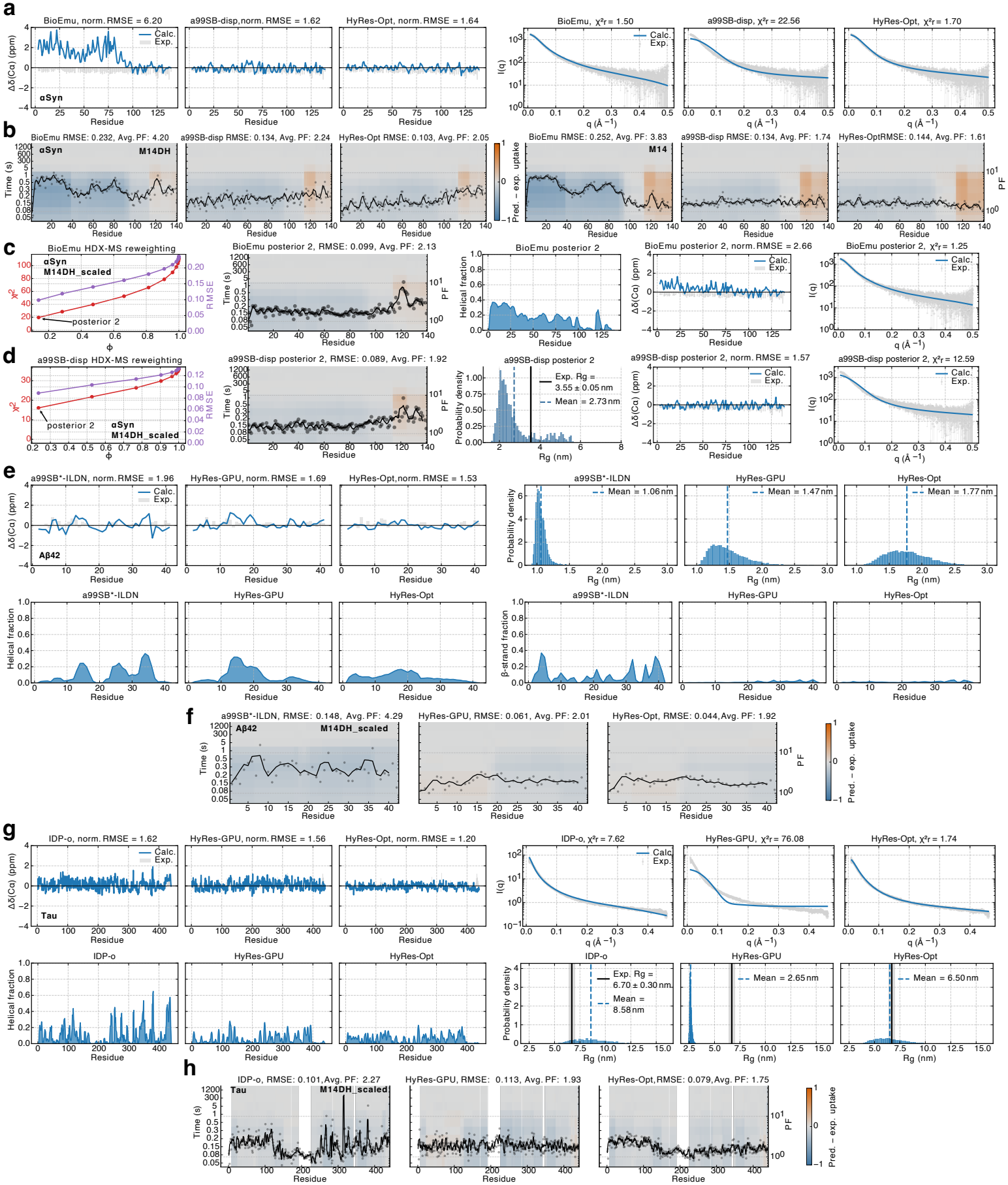
Evaluating and ranking disordered protein ensembles with millisecond HDX-MS. **a**. Comparison of calculated and experimental C*α* chemical shifts (left) and SAXS scattering curves (right) for three different *α*-synuclein ensembles. **b**. Difference between predicted and experimental per-peptide deuterium uptake across time (heatmaps; left y-axis, overall RMSE indicated) and ensemble-averaged PFs (right y-axis with running average over three residues) for the three *α*-synuclein ensembles. Heatmaps show values across the protein sequence, with residues covered by multiple peptides averaged across all overlapping peptides. Two forward model variants are shown to highlight the effect of including electrostatic contributions in the calculation (M14DH). **c**. Left to right: HDX-MS reweighting (with the M14DH_scaled model) of the BioEmu *α*-synuclein ensemble showing agreement with the HDX data as a function of the effective fraction of frames, *ϕ*; difference between predicted and experimental per-peptide deuterium uptake across time; helical propensities after reweighting (posterior 2); agreement with *Cα* chemical shifts and the SAXS scattering curve after reweighting. **d**. Left to right: HDX-MS reweighting of the a99SB-disp *α*-synuclein ensemble showing agreement with the HDX data as a function of the effective fraction of frames, *ϕ*; difference between predicted and experimental per-peptide deuterium uptake across time; R_*g*_ mean and distribution (posterior 2); agreement with *Cα* chemical shifts and the SAXS scattering curve after reweighting. **e**. Analysis of three different A*β*42 ensembles showing a comparison of calculated and experimental C*α* chemical shifts, R_*g*_ distributions and secondary structure propensities. **f**. Difference between predicted and experimental per-peptide deuterium uptake across time (heatmaps; left y-axis, overall RMSE indicated) and ensemble-averaged PFs (right y-axis with running average over three residues) for the three A*β*42 ensembles using the M14DH_scaled model. **g**. Analysis of three different Tau ensembles showing a comparison of calculated and experimental C*α* chemical shifts, SAXS scattering curves, helical structure propensities, and R_*g*_ distributions. **h**. Difference between predicted and experimental per-peptide deuterium uptake across time (heatmaps; left y-axis, overall RMSE indicated) and ensemble-averaged PFs (right y-axis with running average over three residues) for the three Tau ensembles using the M14DH_scaled model.

**Supplementary Figure 9.**
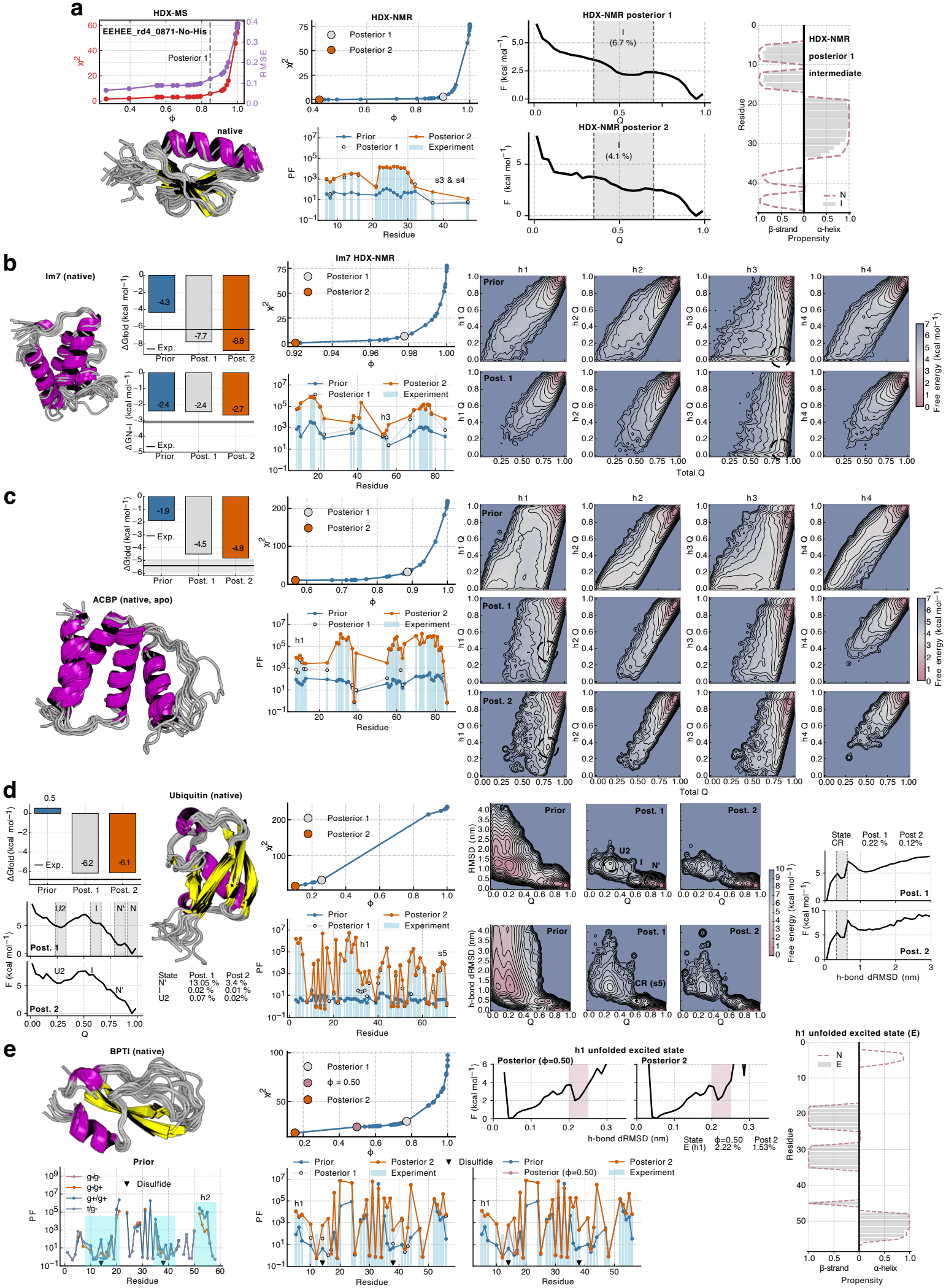
Characterisation of intermediate and excited states by HDX reweighting. See next page for full caption. **a**. Analysis of EEHEE_rd4_0871 after HDX-MS (left) and HDX-NMR (right) reweighting. Experimental and calculated HDX-NMR PFs are compared before and after reweighting; free-energy landscapes are shown as a function of the fraction of native contacts (*Q*), together with secondary structure propensities of the intermediate (I) and native (N) states after HDX-NMR reweighting. **b**. Analysis of Im7 HDX-NMR reweighting. Left to right: experimental and calculated folding free energies, including the free-energy difference between the intermediate (I) and native (N) states; agreement with experimental ln(PF) values as a function of the effective fraction of frames (*ϕ*) and comparison of experimental and calculated PFs; and two-dimensional free-energy landscapes as a function of total-protein *Q* and *Q* for individual helices before and after reweighting. Black circles indicate a local minimum corresponding to the I state with helix h3 unfolded. **c**. Analysis of ACBP HDX-NMR reweighting. Left to right: experimental and calculated folding free energies; agreement with experimental ln(PF) values as a function of *ϕ* and comparison of experimental and calculated PFs; and two-dimensional free-energy landscapes as a function of total-protein *Q* and *Q* for individual helices before and after reweighting. Black circles indicate a local minimum corresponding to the I state with helix h1 partially unfolded. **d**. Analysis of ubiquitin HDX-NMR reweighting. Left to right: experimental and calculated folding free energies and folding landscapes after reweighting (posterior 1 and posterior 2), including populations of identified conformational states; agreement with experimental ln(PF) values as a function of *ϕ* and comparison of experimental and calculated PFs; two-dimensional free-energy landscapes as a function of *Q* and backbone RMSD (top) or *β*-strand s5 hydrogen-bond distance RMSD from the native state (bottom), with black circles indicating identified conformational states; and one-dimensional free-energy landscapes after reweighting as a function of s5 hydrogen-bond distance RMSD, including populations of the C-terminally retracted (CR) state. **e**. Analysis of BPTI HDX-NMR reweighting. Left to right: predicted PFs of BPTI substates with different Cys14–Cys38 disulfide-bond conformations (g, gauche; t, trans), highlighting local differences near the disul-fide bridge and C-terminal helix h2 (shaded regions); agreement with experimental ln(PF) values as a function of *ϕ* and comparison of experimental and calculated PFs for the posterior ensembles indicated on the reweighting L-curve; free-energy landscapes as a function of helix h1 hydrogen-bond distance RMSD from the native state, with populations of the excited (E) state after reweighting; and secondary structure propensities of the h1-unfolded E state compared with the native (N) state.

## Supplementary Data

A supplementary data Excel file containing a summary of the HDX-MS experimental parameters, replication table and sequence coverage is provided.

